# VipariNama: RNA vectors to rapidly reprogram plant morphology and metabolism

**DOI:** 10.1101/2020.06.03.130179

**Authors:** Arjun Khakhar, Cecily Wang, Ryan Swanson, Sydney Stokke, Furva Rizvi, Surbhi Sarup, John Hobbs, Daniel F. Voytas

## Abstract

Synthetic transcription factors have great promise as tools to explore biological processes. By allowing precise alterations in gene expression, they can help elucidate relationships between gene expression and plant morphology or metabolism. However, the years-long timescales, high cost, and technical skill associated with plant transformation have dramatically slowed their use. In this work, we developed a new platform technology called VipariNama (ViN) in which RNA vectors are used to rapidly deploy synthetic transcription factors and reprogram gene expression *in planta*. We demonstrate how ViN vectors can direct activation or repression of multiple genes, systemically and persistently over several weeks, and in multiple plant species. We also show how this transcriptional reprogramming can create predictable changes to metabolic and morphological phenotypes in the model plants *Nicotiana benthamiana* and *Arabidopsis thaliana* in a matter of weeks. Finally, we show how a model of gibberellin signaling can guide ViN vector-based reprogramming to rapidly engineer plant size in both model species as well as the crop *Solanum lycopersicum* (tomato). In summary, using VipariNama accelerates the timeline for generating phenotypes from over a year to just a few weeks, providing an attractive alternative to transgenesis for synthetic transcription factor-enabled hypothesis testing and crop engineering.

## Introduction

There is an urgent need for tools that accelerate the pace of biological discovery in crop plants, so mechanistic insights can be leveraged to enhance crops and help deliver global food security^1,2^. By examining the genomic changes underlying the domestication of wild plants into modern crops, it has become clear that many desirable traits are driven by changes in the expression of certain key genes^3,4^. These insights highlight how mechanistic models that relate gene expression to development and metabolism could be used to determine the specific changes in gene expression required to obtain agriculturally beneficial phenotypes.

Building and validating robust mechanistic models of how gene expression relates to whole plant phenotype relies on the ability to study the phenotypic outcome of controlled changes to gene expression. These changes could be implemented in a single step using synthetic transcription factors with programmable DNA-binding domains, such as Cas9^5^. These tools allow the exploration of intermediate ranges gene expression inaccessible by traditional methods like overexpression or knockouts. Synthetic transcription factors have already been used to make predictable alterations in expression in a range of plants^5–7^. This strategy of model validation was first demonstrated in *A. thaliana*, where a mathematical model of auxin regulated branching was validated through the deployment of a <u>H</u>ormone <u>A</u>ctivated <u>C</u>as9-based <u>R</u>epressor (HACR) to reprogram shoot architecture^5^.

While this work provided a proof-of-concept for deployment of synthetic transcription factors for the elucidation of biological mechanisms, the extension of this strategy into crop plants faces a hurdle shared by most applications of plant synthetic biology: the challenges of generating transgenic plants. These challenges include high costs and technical skill, which restrict access to high resource settings, as well as a sometimes year-long time scales, slowing the iterative designbuild-test cycle of synthetic biology to a crawl^8^. In this work, we sought to circumvent this core challenge with a new platform called VipariNama, the Sanskrit word meaning ‘to change’. VipariNama uses RNA vectors (ViN vectors), which can spread throughout the plant and persist over time, to deliver synthetic transcription factors and make alterations to gene expression, and thereby rapidly engineer phenotypes without necessitating transgenesis. This approach conceptually parallels the use of RNA vectors in gene therapies^9^, but with the end goal of reengineering plant metabolism and morphology to elucidate or validate mechanistic models of biology.

Positive single stranded RNA viruses are an ideal starting point to build ViN vectors from, as they are systemically mobile and persist for long periods of time in plants^10^. Some of these viruses can have very broad host ranges and are largely asymptomatic. For these reasons we decided to base our ViN vectors on the Tobravirus, Tobacco Rattle Virus (TRV)^10^. Previous work has identified regions of the TRV genome into which foreign gene sequences can be added to re-purpose these viruses as protein production tools^11^. However, using the same strategy to deliver synthetic transcription factors, such as the HACR, is not possible due to the limited loading capacity of the virus, namely less than 1kb^12^. Larger cargos abrogate the viruses’ capacity for systemic movement through the plant and are often lost through recombination.

To overcome this challenge, we developed VipariNama 1.0, wherein we created a transgenic plant constitutively expressing a Cas9-based synthetic transcription factor and used ViN vectors to deliver single guide RNAs (sgRNAs) to specify targets for this transcription factor. Next, we demonstrated how this strategy can be made more flexible using sgRNA scaffolds to simultaneously generate activation and repression-based phenotypes in a plant. Finally, we showed how this platform could be made even more flexible by building an ensemble of ViN vectors to deliver nearly all the synthetic transcription factor components to a plant stably expressing only Cas9. This system, VipariNama 2.0, can implement persistent activation or repression of multiple genes across several plant species. We also demonstrated ViN vectors can be used in conjunction with mechanistic models to rapidly engineer metabolic and morphological phenotypes. A model of gibberellin signaling^13^ was used to direct targeting of synthetic transcription factors delivered via ViN vectors try and engineer plant size, an agriculturally important trait, in the model plants *A. thaliana* and *N. benthamiana*, as well as the crop, tomato.

**Graphical abstract.**
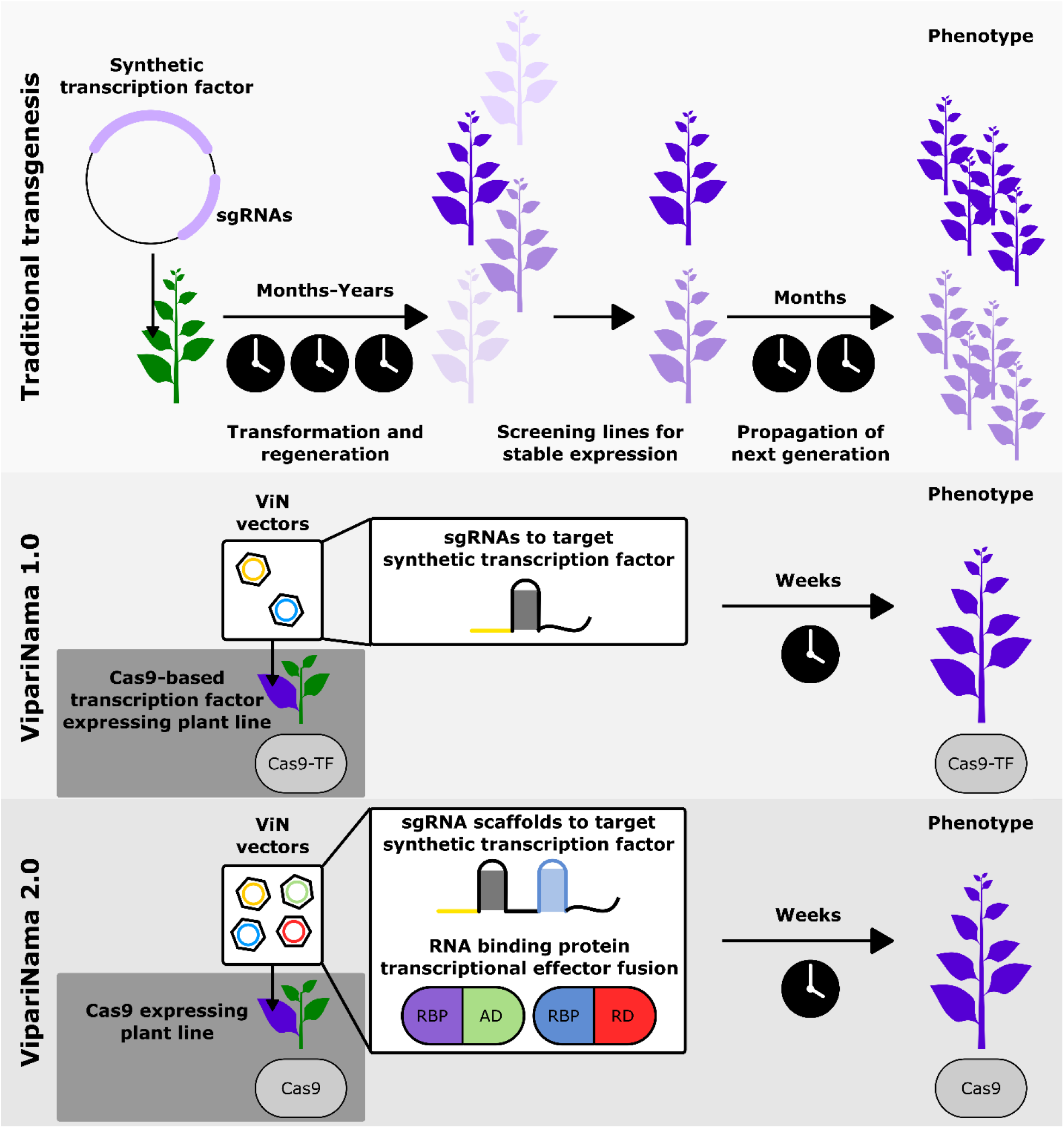
VipariNama allows rapid reprograming of plant morphology and metabolism by delivering synthetic transcription factors on RNA vectors. Schematics depicting how synthetic transcription factors can be deployed to perturb gene expression and create phenotypes via traditional transgenesis (top), VipariNama 1.0 (middle), where ViN vectors deliver sgRNAs to a plant that is stably expressing Cas9-based transcription factors, and VipariNama 2.0 (bottom), where ViN vector ensembles deliver both sgRNA scaffolds as well as RNA binding protein-transcriptional effector fusions.

## Results

### ViN vectors deliver gRNAs to GA-HACR *Nicotiana benthamiana* lines and implement repression

Previous characterization of the GA-HACR, a gibberellin responsive Cas9-based repressor, demonstrated it is an effective tool to study gibberellin (GA) signaling *in planta*^5^. HACRs consist of a deactivated Cas9 protein (dCas9) fused to a phytohormone degron and a transcriptional repression domain. When complexed with a co-expressed sgRNA, a HACR is targeted to repress expression of a specific gene. In the presence of a threshold phytohormone concentration, the HACR is targeted for degradation, thus relieving repression. In lines that have a stably integrated GA-HACR biosensor, a significant increase in luciferase signal is observed in response to external GA treatment or internal GA biosynthesis^5^. As the TRV parent virus for ViN vectors has been previously shown to propagate well in *N. benthamiana*, we created transgenic lines of this species constitutively expressing a GA-HACR and used these plants to test the capacity of ViN vectors to reprogram transcription (Figure 1A). Our aim was to target the GA-HACR to repress the expression of the *GA20ox* genes in the transgenic lines. These genes are responsible for GA biosynthesis, and their GA-dependent expression is an important parameter controlling the concentration of GA in the cell^13^.

**Figure 1.**
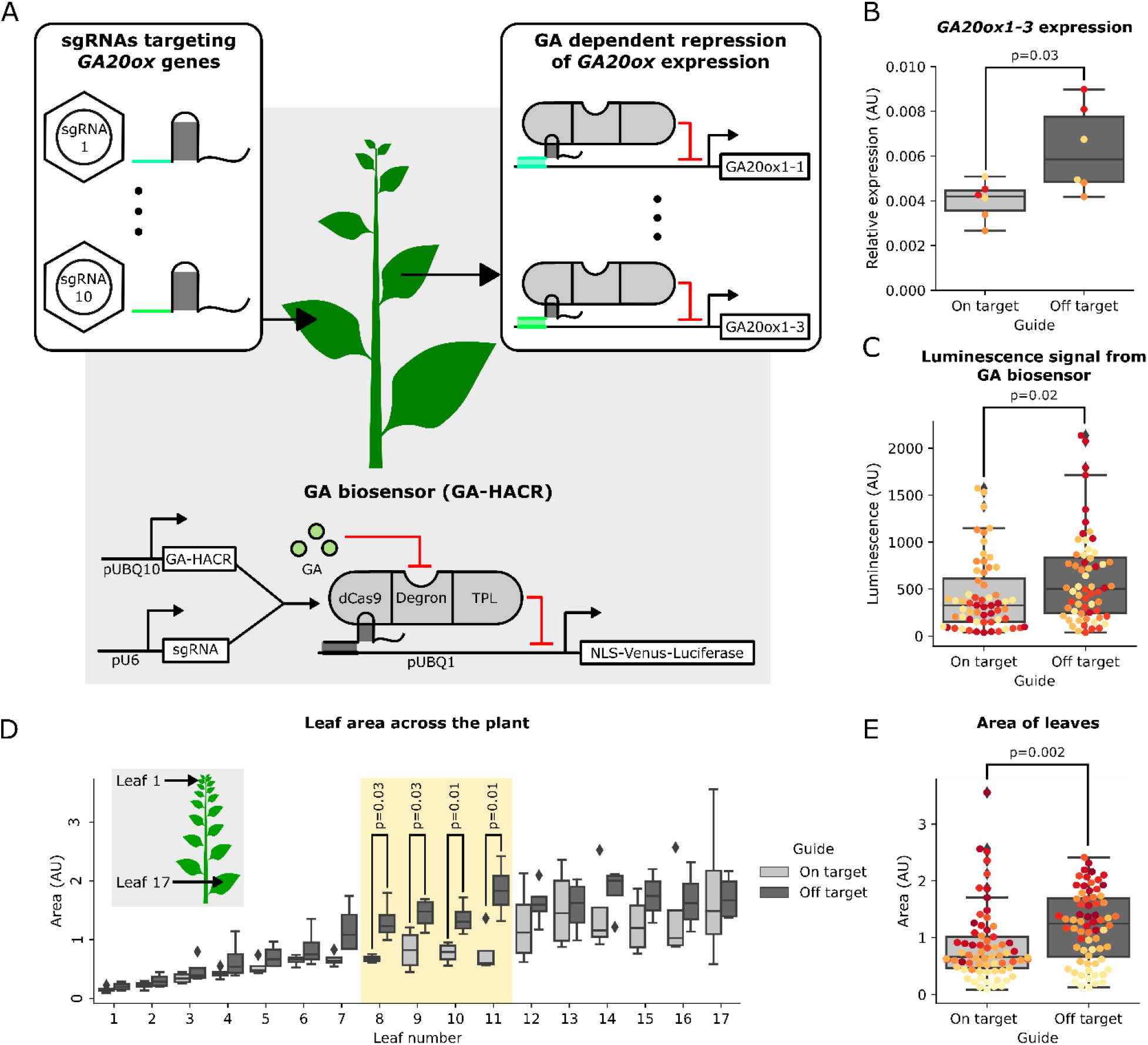
ViN vectors can deliver sgRNAs to reprogram a HACR to repress *GA20ox* expression and create a reduction in GA levels and an associated reduction in leaf area. A) Schematic describing the GA-HACR based GA biosensor that is integrated into the *Nicotiana benthamiana* genome. Here, the constitutively expressed GA-HACR complexes with a sgRNA that targets it to repress a luciferase reporter in a GA-dependent manner. This reporter creates a luminescence signal in response to GA in a dose dependent manner. The inset panels show ViN vectors delivering sgRNAs to target the GA-HACR to the *GA20ox* genes in *N. benthamiana* (left) and resultant reprogrammed HACRs in systemic tissues (right). B) Box plots summarizing relative expression of *GA20ox1-3*, normalized to the *EF1alpha* housekeeping gene, from systemic leaves of the plant lines described in panel A that were treated with ViN vectors encoding on-target (light grey) or off-target (dark grey) sgRNAs. Each dot of the same color represents data from independent biological replicates (n=3 per treatment). Reported p-values were calculated using a t-test. C) Box plots summarizing luminescence signal of the GA biosensor from the leaves of plants treated with ViN vectors encoding on-target (light grey) or off-target (dark grey) sgRNAs. Each dot of the same color represents data from leaves on an independent biological replicate (n=4 per treatment). Reported p-values were calculated using a t-test. D) Box plots summarizing the area of leaves of the plants described in panel A treated with ViN vectors encoding on-target (light grey) or off-target (dark grey) sgRNAs. The inset shows all the leaves assayed across the plant, with leaf number 1 being the top leaf. The yellow region highlights the leaves in which a statistically significant difference in size between plants treated with on-target and off-target sgRNAs was observed. E) Box plots summarizing the area of all the leaves plotted together. Each dot represents data from a leaf of an independent biological replicate (n=4 per treatment). Reported p-values were calculated using a t-test.

ViN vectors were built to encode sgRNAs targeting two sites within the first 500 base pairs upstream of the transcription start sites of five putative *GA20ox* genes. These vectors were delivered via Agrobacterium infiltration to young plants in parallel with control vectors that encoded sgRNAs with no specific targets in the *N. benthamiana* genome. We used this same strategy for on- and off-target sgRNA design, as well as vector delivery, in all the other experiments described in this work.

RNA was extracted from tissue from the fourth leaf above the infiltrated leaf (the 4^th^ systemic leaf), twenty-four days after vector delivery and expression of the target genes was quantified with qRT-PCR. We observed significant repression of the expression of the *GA20ox1-3* gene in plants treated with ViN vectors encoding the on-target sgRNAs compared to plants treated with the off-target control (Figure 1B). We also observed a decrease in the median expression of the two more highly expressed *GA20ox* genes targeted, *GA20ox1-1* and *GA20ox1-2*, however the effect was not significant (Figure 1 Supplementary Figure 1). This might be because these strongly expressed genes are more challenging to repress^6^. The other two GA20ox genes we targeted, *GA20ox1D-1* and *GA20ox1D-2*, were not expressed in the leaves at significant levels and so we would not expect to observe repression (Figure 1 Supplementary Figure 1). These results demonstrate how ViN vectors can be applied to reprogram synthetic Cas9-based transcription factors by delivering the appropriate sgRNAs. These results also show that the transcriptional regulation conferred by this approach can be observed in systemic leaves for several weeks postdelivery.

### ViN vector-mediated transcriptional reprogramming in GA-HACR plant lines generates metabolic and developmental phenotypes

The GA-HACR line we built incorporates the previously described GA-biosensor (Figure 1A). As the GA-HACR is targeted to regulate a luciferase reporter, also included in this construct, in a GA dependent manner, treatment of these plants with luciferin substrate should result in a luciferase signal proportional to the concentration of GA within that tissue^5^. We used this reporter to compare GA levels in GA-HACR *N. benthamiana* plants to which we had delivered sgRNAs targeting the *GA20ox* gene family or off-target controls. Based on a mathematical model of the GA signaling pathway, increasing the GA-dependent repression of the GA biosynthesis genes, the *GA20ox* gene family, should result in a decrease in cellular GA concentration^13^. We observed a significant decrease in the luminescence observed from plants treated with on-target sgRNAs compared to the controls (Figure 1C). This is consistent with the model predicting that a decrease in GA-dependent *GA20ox* expression should lower GA levels.

Reduction in GA levels should also result in an increased accumulation of *DELLA* proteins, as GA triggers their proteasomal degradation, and an associated dwarfing in leaf tissue^14–16^. We phenotyped plants treated with ViN vectors five weeks post infection and observed a significant decrease in leaf area in plants treated with on-target sgRNAs compared to plants treated with control sgRNAs (Figure 1E). This effect was particularly pronounced in the leaves closer to the infiltration site (leaf numbers 8-11) (Figure 1D). We hypothesize this was because these leaves were infected early in development, unlike older leaves, and had sufficient time to fully expand by the time of measurement, in contrast to younger, still-developing leaves. These results were replicated in an independently grown set of plants, confirming that they were not due to growth conditions or relative plant health (Figure 1 Supplementary Figure 2). Taken together these results serve as a proof-of-concept of how ViN vectors can be deployed in HACR lines to alter transcriptional landscapes and rapidly reprogram metabolism, i.e. GA biosynthesis, as well as morphology, i.e. leaf area.

**Figure 1, Supplementary Figure 1.**
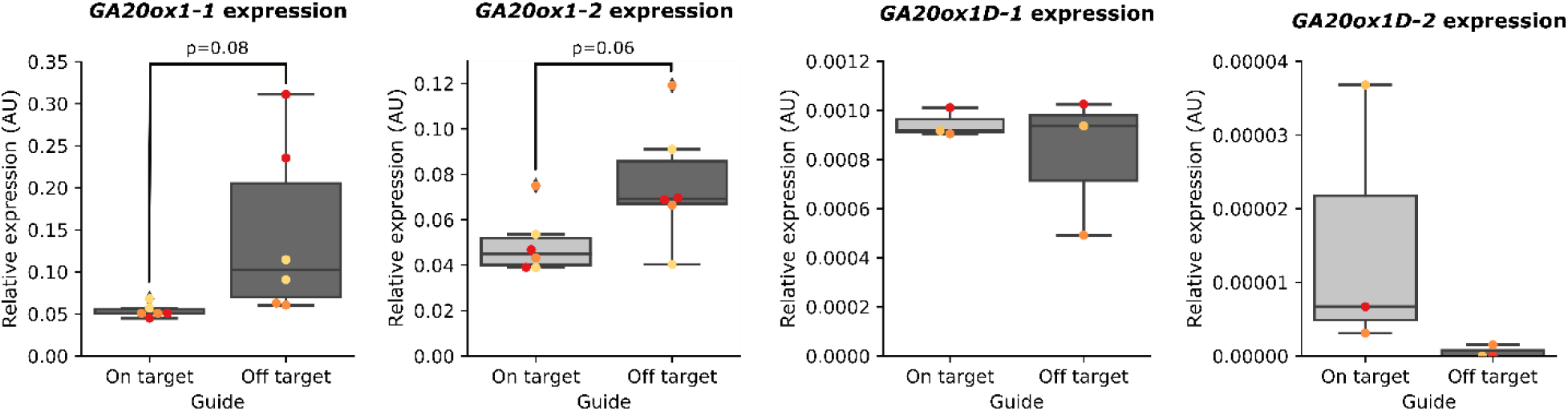
Relative expression of the other GA20ox genes targeted in on and off-target plants. Box plots summarizing expression of *GA20ox1-1, GA20ox1-2, GA20ox1D-1*, and *GA20ox1D-2*, normalized to the *EF1alpha* housekeeping gene, from systemic tissue of the plant lines described in panel A of Figure 1 that were treated with ViN vectors encoding on-target (light grey) or off-target (dark grey) sgRNAs. Each dot of the same color represents data from independent biological replicates (n=3 per treatment). Reported p-values were calculated using a t-test.

**Figure 1, Supplementary Figure 2.**
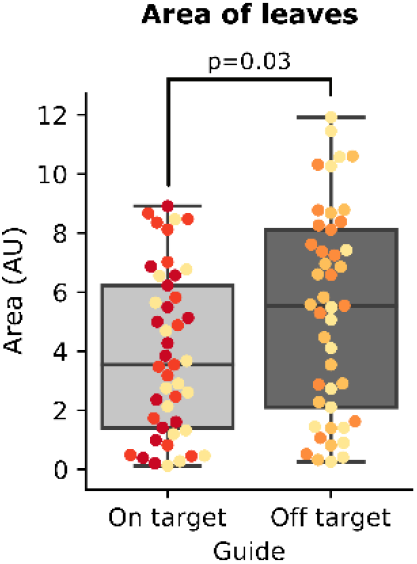
Phenotype alterations created by ViN 1.0 in *N. benthamiana* GA-HACR lines can be replicated in a second set of independently grown plants. Box plots summarizing the area of all the leaves from plants treated with ViN vectors encoding on-target (light grey) or off-target (dark grey) sgRNAs. Each dot of the same color represents data from a leaf of an independent biological replicate (n=3 per treatment). Reported p-values were calculated using a t-test.

### sgRNA scaffolds delivered via ViN vectors simultaneously create activation and repression phenotypes

One potential drawback of using HACR lines for reprogramming plant phenotypes is that certain traits require phytohormone independent regulation, or simultaneous activation and repression of multiple genes with different effectors. This would require an orthogonal dCas9 variant for each effector and the construction of a new transgenic line for each unique combination, thereby slowing down hypothesis testing. To overcome this limitation, we designed plant lines that enable scaffold-based reconstitution of transcription factors. Multiple constitutive expression cassettes are incorporated into these plant’s genome: a Cas9 cassette to provide programable DNA binding and cutting, and a set of unique RNA binding proteins fused to different transcriptional effectors (Figure 2A). ViN vectors can then be used to deliver specialized sgRNA scaffolds to these plants, which serve to direct Cas9 to a genomic target, but also to recruit specific transcriptional effectors. By truncating the target site of the sgRNA to 14 base pairs in these scaffolds, the nuclease-active Cas9 is directed to bind to DNA, but not cut it^17^. The addition of specific motifs at the 3’ end of the scaffold enables interaction with a specific RNA binding protein fused to a transcriptional effector. Together, these components reconstitute the desired transcription factor at the locus of interest^18^ (Figure 2A). In principle, this would enable simultaneous activation and repression of different genes *in planta*. Delivering a full-length sgRNA, rather than a truncated sgRNA scaffold, to these same lines would allow targeted gene ablation, because the Cas9 nuclease is active. This strategy has been demonstrated previously for efficient somatic genome editing^19^.

**Figure 2.**
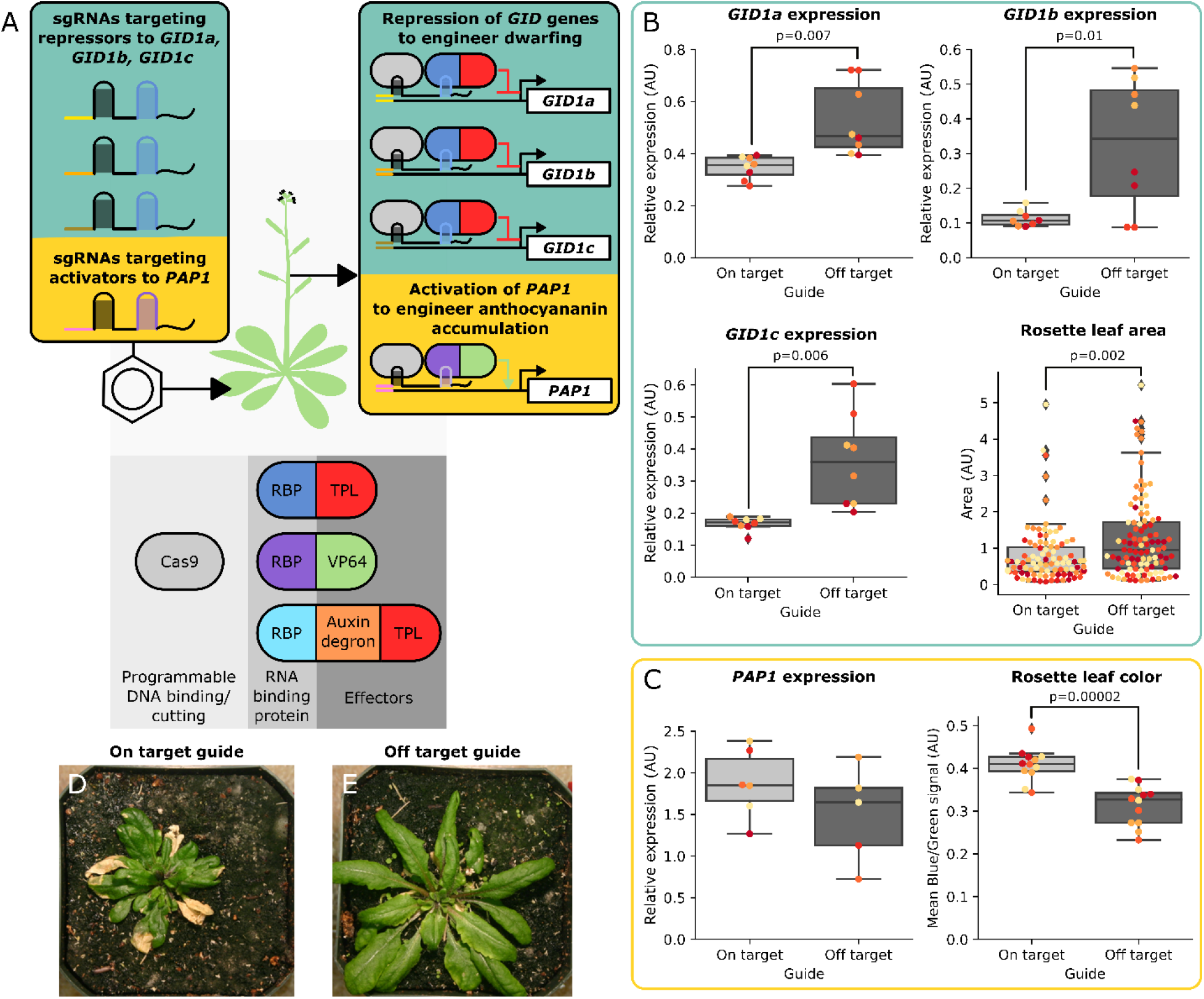
ViN vectors deliver sgRNA scaffolds to plant lines that enable scaffold-based reconstitution of transcription factors to implement transcriptional regulation and alter phenotypes. A) Schematic describing components expressed by an *A. thaliana* line to enable scaffold-based reconstitution of transcription factors. This line constitutively expresses nuclease active Cas9 as well as three RNA binding proteins fused to either a truncation of the *TOPLESS* repressor from *A. thaliana*, the VP64 activator, or an auxin degron which is also fused to a *TOPLESS* repressor. The colored insets describe the sgRNA scaffolds being delivered to this line via ViN vectors (left) and the resultant transcriptional perturbations as well as the expected phenotypic results (right). B) Box plots summarizing the expression of the *GID1a*, *GID1b* and *GID1c* genes, normalized to the *PP2A* housekeeping gene, in rosette leaves as well as the rosette leaf area of the plants described in panel A that were treated with ViN vectors encoding on-target (light grey) or off-target (dark grey) sgRNAs. Each dot of the same color represents data from independent biological replicates (n=4 per treatment). Reported p-values were calculated using a t-test. C) Box plots summarizing data collected from the rosette leaves of plants described in panel A that were treated with ViN vectors encoding on-target (light grey) or off-target (dark grey) sgRNAs three weeks after delivery. The box plot on the left summarizes the expression of the *PAP1* gene, normalized to the *PP2A* housekeeping gene, in rosette leaves. The box plot on the right summarizes the average blue signal normalized to the green signal, which is an established proxy for anthocyanin concentration^23^, from images of rosette leaves. Each dot of the same color represents data from leaves of independent biological replicates (n=5 per treatment for expression and n=4 per treatment for leaf color). Reported p-values were calculated using a t-test. D,E) Representative pictures of rosettes of plants treated with on-target (D) or off-target (E) sgRNAs at the time of phenotyping.

To quickly test this system, we built lines of *A. thaliana* encoding the previously described components (Figure 2A). ViN vectors were then used to simultaneously deliver a set of sgRNA scaffolds targeting a repressor to the three *GID* genes, which are GA receptors^13^, and an activator to the *PAP1* gene, which is a *MYB* transcription factor^20,21^ (Figure 2A). Tissue was collected from systemic rosette leaves 24 days postdelivery. When we compared expression of the three *GID* genes in plants treated with on-target sgRNAs to controls we see a significant repression of all three genes (Figure 2B). Previous studies and the model of GA signaling both suggest that reducing *GID* expression should result in dwarfing, due to the increased concentration of growth-inhibiting *DELLA* proteins^13,22^. We observed a significant reduction of rosette leaf area in plants treated with on-target sgRNAs compared to plants treated with off-target controls (Figure 2B).

We do not observe a statistically significant increase in the expression of the *PAP1* gene in plants treated with the on-target sgRNAs compared to controls (Figure 2C). This might be because the gene is natively highly expressed: *PAP1* expression in control plants is five times the *GID* expression levels, making any increase in expression subtle at best^6^. However, as *PAP1* is a master regulator of the anthocyanin biosynthesis pathway, even subtle changes in expression could result in a phenotype. Previous studies also show that overexpression of *PAP1* results in anthocyanin accumulation^20,21^. The average ratio of blue to green signal in images of leaves is an established proxy for anthocyanin accumulation^23^. We observed a significant increase in the average ratio of blue to green signal in images of rosette leaves of plants treated with on-target sgRNAs compared to controls (Figure 2C). These results imply that while the increase in *PAP1* expression is too subtle to see in the leaves assayed, it may still be capable of producing the increased accumulation of purple anthocyanin pigment. To demonstrate that these were two separate phenotypes, and not just two phenotypes caused by the observed *GID* repression, we repeated the experiment in a plant line without the repressor targeted to the *GID* genes. We again observed a significant increase in the average ratio of blue to green signal in images of rosette leaves of plants treated with on-target sgRNAs, consistent with anthocyanin accumulation from increased PAP1 expression (Figure 2 Supplementary Figure 1). This demonstrates that these are independent phenotypes. Taken together these results show how ViN vectors can be used in conjunction with plant lines that enable scaffold-based reconstitution of transcription factors to reprogram regulation and simultaneously create morphological and metabolic phenotypes within weeks.

**Figure 2 Supplement 1.**
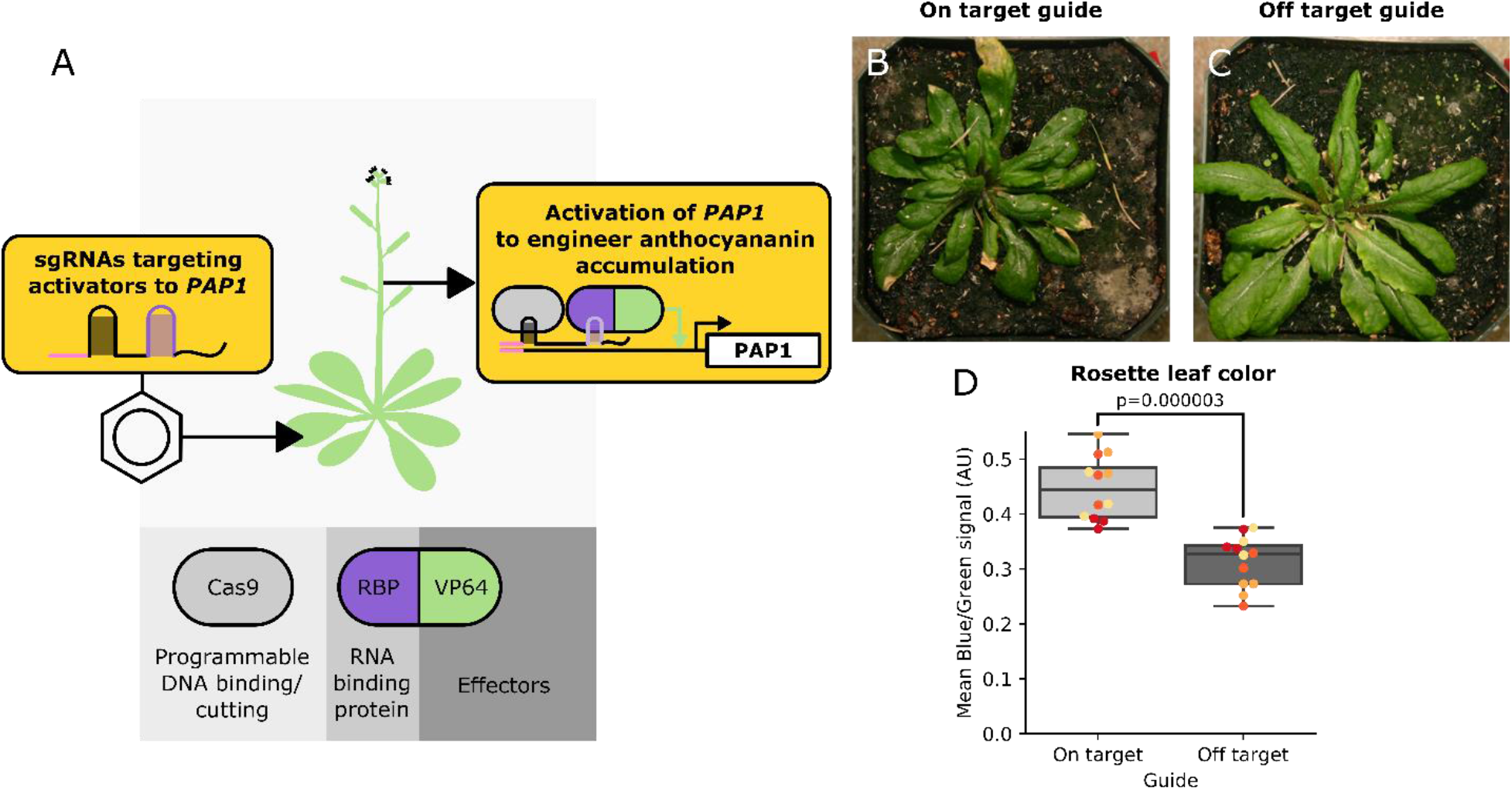
*PAP1*-associated anthocyanin phenotype can be obtained independently of GID repression. A) Schematic describing an *A. thaliana* line engineered to enable scaffold-based reconstitution of transcription factors. This line constitutively expresses nuclease active Cas9 as well as an RNA binding protein fused to the VP64 activator. The colored insets describe the sgRNA scaffolds being delivered to this line via ViN vectors (left) and the resultant transcriptional perturbation as well as the expected phenotypic result (right). B,C) Representative pictures of rosettes of plants treated with on-target (B) or off-target (C) sgRNAs at the time of phenotyping. D) Box plot summarizing the average blue signal normalized to the green signal, which is an established proxy for anthocyanin concentration^23^, from images of rosette leaves of the plants described in panel A that were treated with ViN vectors encoding on-target (light grey) or off-target (dark grey) sgRNAs. Each dot of the same color represents data from leaves of independent biological replicates (n=3 per treatment). Reported p-values were calculated using a t-test.

### ViN vector ensembles can implement transcriptional regulation in Cas9 expressing plant lines

While plant lines engineered to enable scaffold-based reconstitution of transcription factors enable flexible transcriptional reprogramming in a plant, this strategy is still limited to the transcriptional effector domains integrated into the genome. It has been shown that transcriptional effector domains can have variable efficacy depending on the locus being targeted^7^, which means in certain cases it might be necessary to screen several domains to achieve the desired effect. This would require generating a new transgenic line each time. To overcome this issue, we envisaged VipariNama 2.0, where ensembles of ViN vectors are used to deliver a combination of sgRNA scaffolds and RNA binding protein-transcriptional effector fusions to a transgenic plant line constitutively expressing Cas9 (Figure 3A). This would enable the reconstitution of the desired transcription factors, in a similar fashion to the *Arabidopsis* lines described previously, however in this case the plant would only need to be expressing Cas9.

**Figure 3.**
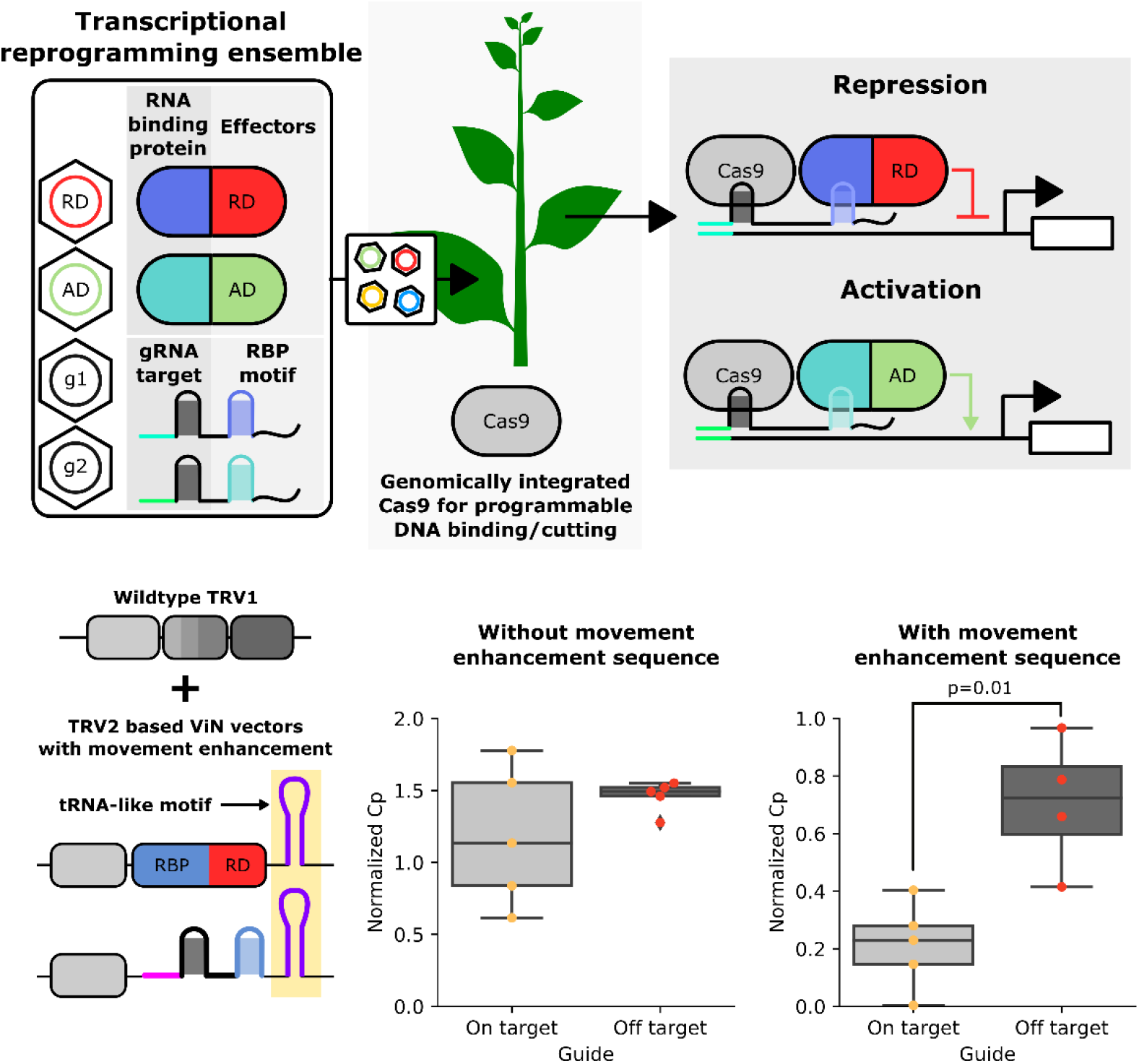
ViN 2.0 vector ensembles composed of vectors with movement enhancement sequences enable transcriptional reprograming in Cas9-expressing plants. A) Schematic describing how VipariNama 2.0 ensembles can be used to reprogram transcription. A ViN 2.0 ensemble consists of a combination of vectors which encode either RNA binding protein-transcriptional effector domain fusions or sgRNA scaffolds (right inset). When these ensembles are delivered to the Cas9 line (center) the components reconstitute the desired transcription factors at the locus of interest (right). B) Schematic describing how the TRV-based vectors in a ViN 2.0 ensemble can be engineered to incorporate movement enhancement motifs (highlighted in yellow). C) Boxplots summarizing expression of *PDS1*, normalized to the expression of the *EF1alpha* housekeeping gene, collected from systemic leaves of Cas9-expressing *N. benthamiana* treated with ViN 2.0 repressor ensembles that encode on-target (light grey) or off-target (dark grey) sgRNA scaffolds. The vectors used in the experiments described in the plot on the left did not contain movement enhancement sequences while the vectors described in the plot on the right did. Each dot represents data from independent biological replicates (n=4-5 per treatment). Reported p-values were calculated using a t-test.

In contrast to the approach of using ViN vectors to deliver just sgRNA scaffolds to a line already expressing a synthetic transcription factor (ViN 1.0), in ViN 2.0 two vectors need to co-localize in the same cell to enable effective transcriptional regulation: the vector encoding the transcriptional effector and the vector encoding the sgRNA scaffold. This is challenging because TRV, like most RNA viruses, tends to move through plants in a non-uniform manner, making cellular co-localization of multiple vectors likely a relatively rare event^24^. We reasoned that colocalization might be improved by incorporating a previously characterized RNA movement enhancement motif into the ViN vectors (Figure 3B). This motif was the first 102 base pairs of the *A. thaliana FLOWERING LOCUS T* (FT) mRNA which adopts a tRNA-like structure^25^. It has been previously shown to be systemically mobile^26,27^ and enhance viral movement^25^.

We built ViN vector ensembles that encoded a repressor, SRDX^7^, as well as sgRNA scaffolds to targeting it to the *PDS1* gene in a Cas9-expressing transgenic line of *N. benthamiana*. One ensemble was built with vectors that had the movement enhancement sequence, and the other without. When we assayed *PDS1* expression in systemic leaves three weeks after vector delivery, we observed significant repression of *PDS1* expression compared to off-target controls in plants treated with the ensemble that included the movement enhancement sequence (Figure 3C). In contrast, the ensemble that did not contain a movement enhancement sequence did not show a significant decrease in expression of *PDS1* (Figure 3C). This is consistent with the hypothesis that greater vector mobility improves efficacy of transcriptional regulation.

We also assayed the vector copy number within the collected tissue using qRT-PCR and observed similar levels in plants treated with ensembles with and without the movement enhancement sequences (Figure 3 Supplementary Figure 1). This further reinforces the idea that the improved regulation is due to an enhanced movement of the vectors rather than an increase in vector concentration due to enhanced stability conferred by the tRNA-like sequence. We also explored if targeting Cas9 to the promoter of the *PDS1* was sufficient to confer repression on its own, which would imply that co-localization of the repressor and sgRNA scaffold was not necessary. We observed that plants that were treated with ensembles that do not encode a repressor showed levels of *PDS1* expression indistinguishable from off-target controls in systemic leaves at three weeks post-delivery (Figure 3 Supplementary Figure 2). This demonstrates that the entire ensemble is necessary for effective regulation. Taken together these results show that ensembles of ViN vectors with movement enhancement sequences can be used to implement systemic transcriptional changes in a Cas9-expressing plant line.

**Figure 3 Supplementary Figure 1.**
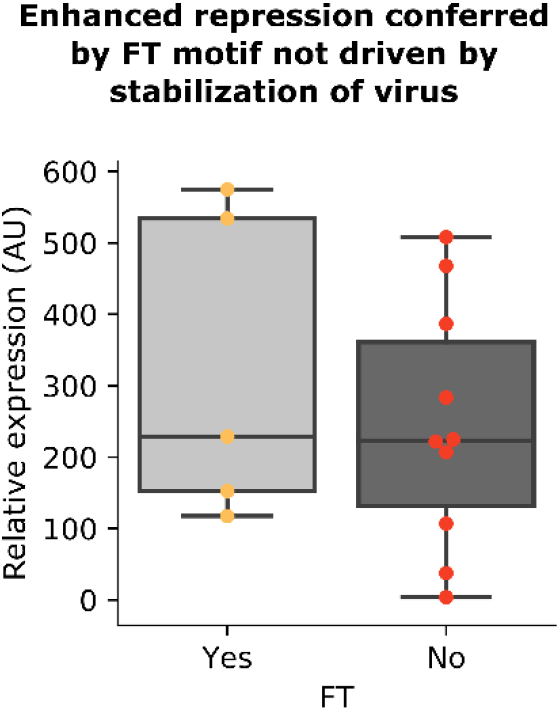
Movement enhancement motif does not confer enhanced stability to the ViN vectors. Boxplots summarizing levels of the ViN vector transcripts, normalized to the expression of the *EF1alpha* housekeeping gene, collected from systemic leaves of Cas9-expressing *N. benthamiana* treated with ViN 2.0 repressor ensembles that either had (light grey) or did not have (dark grey) the tRNA-like motif from the *A. thaliana* FT transcript. Each dot represents data from independent biological replicates (n=5-10 per treatment).

**Figure 3 Supplementary Figure 2.**
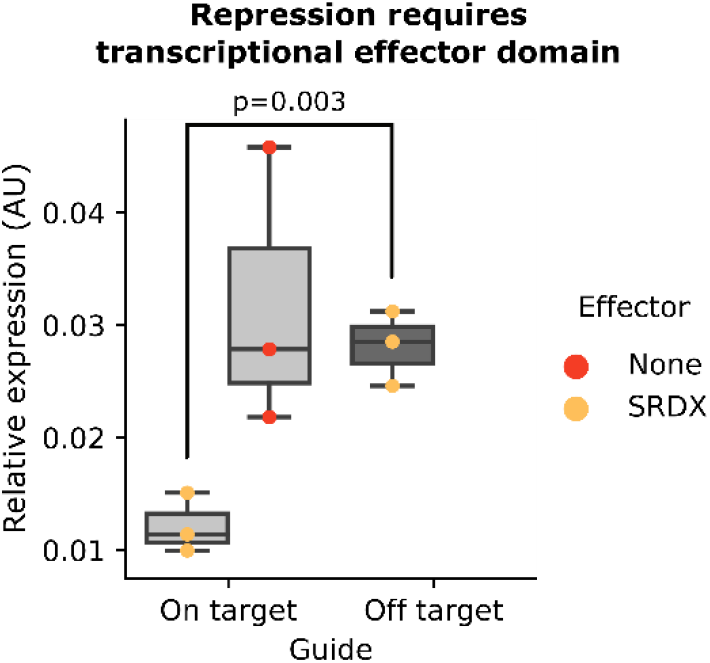
ViN 2.0 based repression requires the presence of the repressor domain. Boxplots summarizing expression of *PDS1*, normalized to the expression of the housekeeping gene *EF1alpha*, collected from systemic leaves of Cas9-expressing *N. benthamiana* treated with ViN 2.0 ensembles that encode on-target (light grey) or off-target (dark grey) sgRNA scaffolds. The red dots represent measurements from plants treated with ensembles that did not encode a repressor, and the yellow dots from plants treated with ensembles that encoded an SRDX repressor. Each dot represents data from independent biological replicates (n=3 per treatment). Reported p-values were calculated using a t-test.

### ViN 2.0 enables easy swapping of effectors and targets

One of the key advantages of ViN 2.0 over ViN 1.0 is the capacity to easily swap effectors by changing the vectors within an ensemble. To demonstrate this, we built an ensemble of ViN vectors, one encoding a VP64 activator and the other encoding sgRNA scaffolds to target the *DFR* gene, a metabolic gene previously described as a good target for activation^6^, in a *N. benthamiana* plant stably expressing Cas9 (Figure 4A). We then collected tissue from systemic leaves three weeks after delivery and quantified the expression of the *DFR* gene using qRT-PCR. We observed a significant increase in the expression of *DFR* in plants treated with an ensemble that encoded on-target sgRNAs compared to plants treated with ensembles which encoded off-target control sgRNAs (Figure 4B).

**Figure 4.**
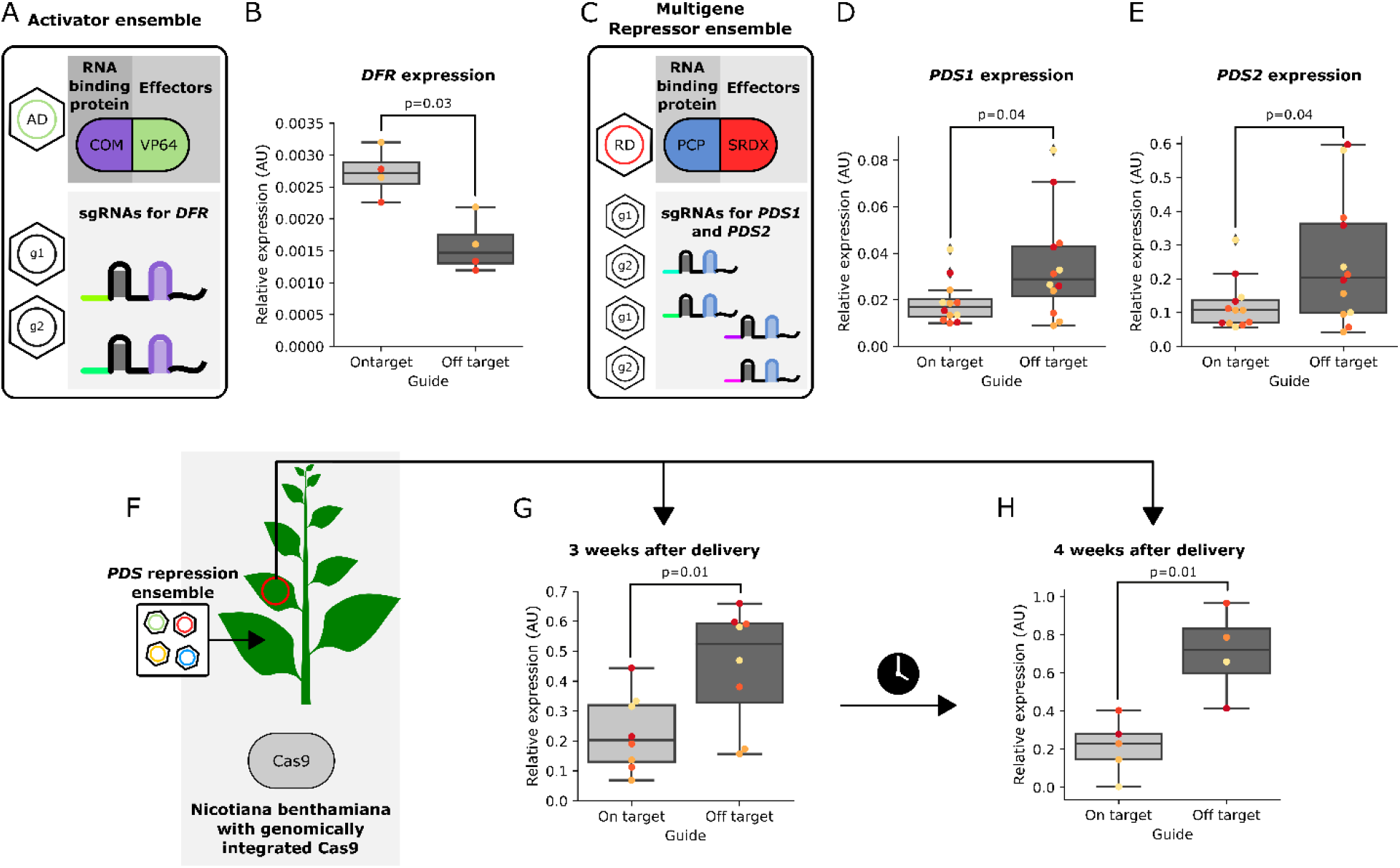
ViN 2.0 ensembles can implement multiplexed repression and activation, systemically and persistently, in a Cas9 expressing plant. A) Schematic depicting the components of the ViN 2.0 activator ensemble used to target the VP64 activator to the *DFR* gene in a *N. benthamiana* line constitutively expressing Cas9. B) Boxplots summarizing normalized *DFR* expression data from the systemic leaves of plants treated with ensembles that encode either on-target (light grey) or off-target (dark grey) sgRNA scaffolds. Each dot of a different color represents data collected from different leaves of independent biological replicates (n = 2 per treatment). Reported p-values were calculated using a t-test. C) Schematic depicting the components of the ViN 2.0 repressor ensemble used to target the SRDX repressor to the *PDS1* and *PDS2* genes in the same line. D,E) Boxplots summarizing *PDS1* (D) and *PDS2* (E) expression data, normalized to the expression of the *EF1alpha* housekeeping gene, from systemic leaves of plants treated with ensembles that encode either on-target (light grey) or off-target (dark grey) sgRNA scaffolds. Each dot of a different color represents data collected from systemic leaves of independent biological replicates (n = 4 per treatment). Reported p-values were calculated using a t-test. F) Schematic depicting a ViN 2.0 repressor ensemble targeted to the *PDS1* gene being delivered to a Cas9-expressing line of *N. benthamiana*, with the 2^nd^ systemic leaf highlighted with a red circle. G,H) Box plots summarizing normalized *PDS1* expression data from the 2^nd^ systemic leaves 3 weeks (G) and 4 weeks (H) after vector delivery from plants treated with ensembles that encode either on-target (light grey) or off-target (dark grey) sgRNA scaffolds. Each dot of a different color represents data collected from systemic leaves of independent biological replicates (n = 4 per treatment). Reported p-values were calculated using a t-test.

Creating certain traits may require the regulation of multiple genes simultaneously due to redundancy, a common phenomenon in plants, which often have duplicated genomes. We set out to test if ViN 2.0 ensembles could be used to alter the regulation of multiple genes simultaneously. We built a new ensemble where one ViN vector encoded an SRDX repressor and the remaining vectors encoded sgRNA scaffolds that target the *PDS1* and *PDS2* genes in a Cas9-expressing *N. benthamiana* line (Figure 4C). We then collected tissue from systemic leaves three weeks after delivery and quantified the expression of *PDS1* and *PDS2* using qRT-PCR. We observed significant repression of expression of both *PDS1* and *PDS2* genes compared to plants treated with ensembles that encode off-target controls (Figure 4D,E). Taken together these results demonstrate how of ViN 2.0 vector ensembles enhance the flexibility of VipariNama based reprograming of transcriptional landscapes *in planta*.

### Characterizing the spatio-temporal gene expression changes conferred by ViN 2.0 ensembles

An important consideration for any mobile vector is characterizing the spatio-temporal trajectory of gene expression through the plant post-delivery. We delivered ensembles targeting a repressor to the *PDS1* gene in a Cas9 expressing *N. benthamiana* line and characterized expression of *PDS1* compared to off-target controls in systemic leaves (Figure 4F). We observed significant repression of *PDS1* expression in the 2^nd^ systemic leaf (two leaves above the leaf delivered to) three weeks after delivery (Figure 4G). We characterized the expression of *PDS1* over time and observed that the repression of expression was still observed at four weeks post vector delivery (Figure 4H).

We also characterized vector abundance throughout the plant post-delivery and observed the expected gradient of vector concentration decreasing from the point of delivery^24^ at two weeks post-delivery (Figure 4 Supplementary Figure 1B). This gradient had equalized by four weeks, with equivalent levels of vector in the second and fourth systemic leaves (Figure 4 Supplementary Figure 1D). We also observed that while we do see the expected trend of decrease in median *PDS1* expression compared to control levels in the 4^th^ and 6^th^ systemic leaves, the difference compared to the off-target control is too small to be significant (Figure 4 Supplementary Figure 2B). This is because the baseline expression of *PDS1* in younger leaves is lower, as can be seen from the progressively lower *PDS1* expression in off-target controls in the 4^th^ and 6^th^ systemic leaves while the on-target expression remains at the same maximally repressed level seen in the 2^nd^ systemic leaf. These results suggest that repression is being implemented systemically. Taken together these results demonstrate how ViN 2.0 vector ensembles can implement systemic and persistent changes to gene expression.

**Figure 4 Supplementary Figure 1.**
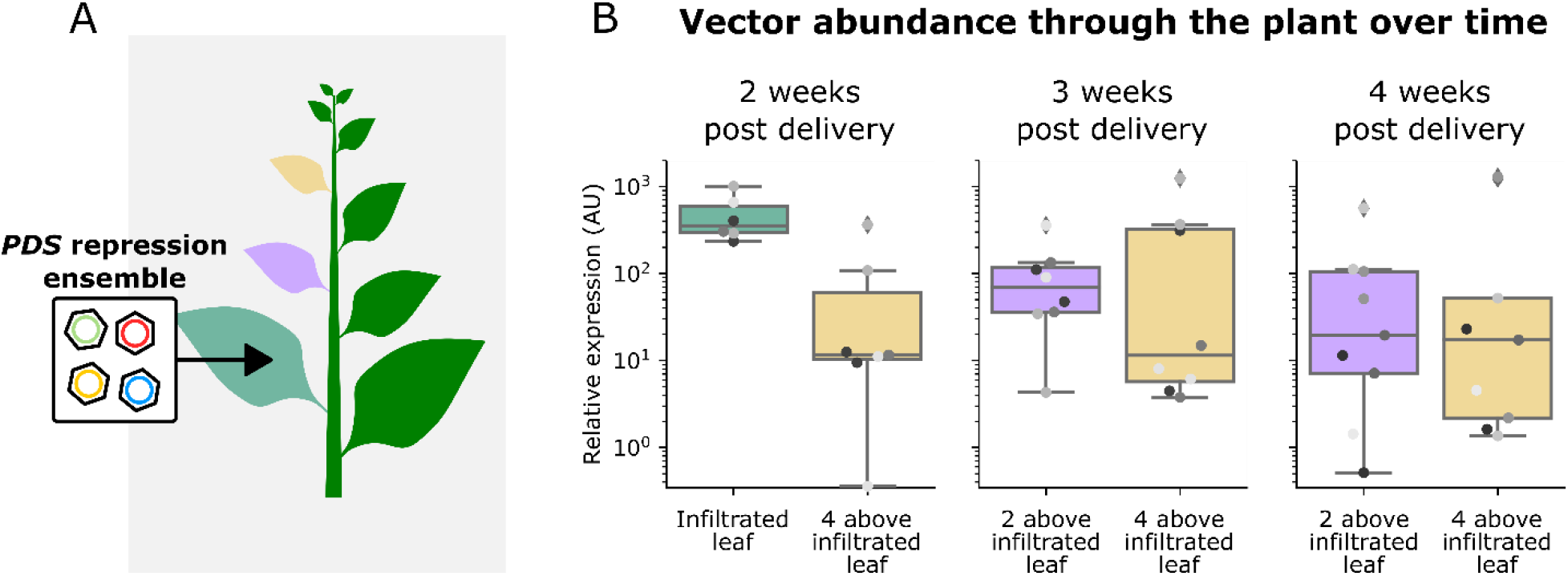
Spatio-temporal quantification of relative ViN vector abundance in the plant. A) Schematic depicting a ViN 2.0 repressor ensemble targeted to the *PDS1* gene being delivered to a Cas9-expressing line of *N. benthamiana*, with different systemic leaves highlighted in non-green colors. B) Box plots summarizing normalized ViN vector RNA levels across the plant at 2 weeks (B), 3 weeks (C), and 4 weeks (D) post vector delivery. Each boxplot is colored to match the color of the leaf it was harvested from in panel A. Each dot represents data collected from systemic leaves of independent biological replicates.

**Figure 4 Supplementary Figure 2.**
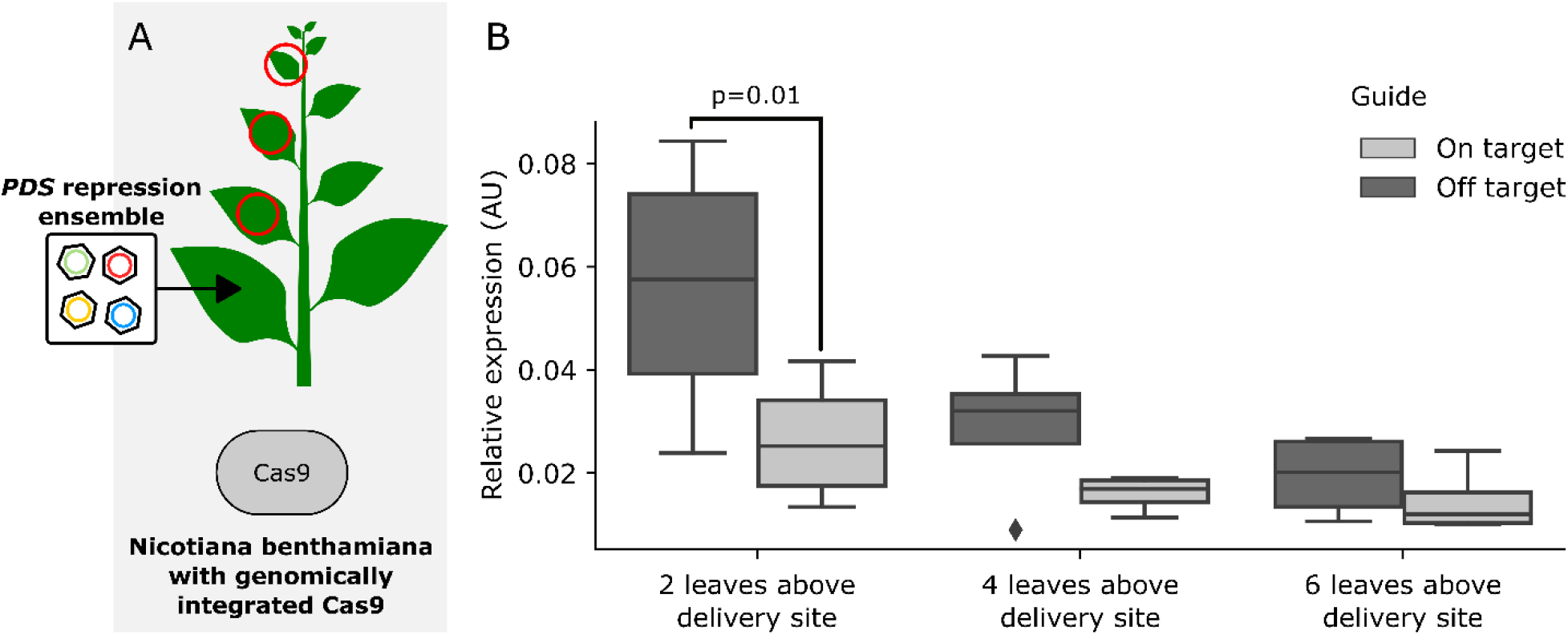
Quantification of *PDS1* repression in tissue progressively distal from the point of infiltration. A) Schematic depicting a ViN 2.0 repressor ensemble targeted to the *PDS1* gene being delivered to a Cas9-expressing line of *N. benthamiana*, with the 2^nd^, 4^th^, and 6^th^ systemic leaves highlighted with a red circle. B) Box plots summarizing normalized *PDS1* expression data, relative to the expression of the *EF1alpha* housekeeping gene, from systemic leaves highlighted in A after 3 weeks post treatment with ensembles that encode either on-target (light grey) or off-target (dark grey) sgRNA scaffolds. Each plot represents data collected from systemic leaves of independent biological replicates (n = 4 per treatment). Reported p-values were calculated using a t-test.

### ViN 2.0 ensembles can rapidly generate transcriptional alternations and associated phenotypic changes in tomato

Our results so far demonstrate the utility of VipariNama to enable rapid hypothesis testing in two model species, *A. thaliana* and *N. benthamiana*. We next set out to test if these vectors could be extended to a crop plant, *Solanum lycopersicum* (tomato). Given our success with creating predictable morphological alterations with ViN 1.0 in *A. thaliana* and *N. benthamiana* through model-guided changes to the GA signaling pathway, we explored if this approach could be applied to alter the stature of tomato. We built ensembles that encode the 188 N-terminal residues of the *TOPLESS* co-repressor from *A. thaliana* and sgRNA scaffolds targeting the tomato *DELLA* protein, *PROCERA* (Figure 5A). This truncation has been previously shown to be sufficient to confer repression^28^ and was necessary to reduce the size of transcriptional effector cargo within the viral cargo capacity. We decided to target *PROCERA*, as this gene has been well studied in tomato^29–31^, and so we could make good predictions of the phenotypic results of altered expression. Additionally, tomato only has a single *DELLA* gene, so targeting it reduces the chance of not observing a phenotype due to genetic redundancy.

**Figure 5.**
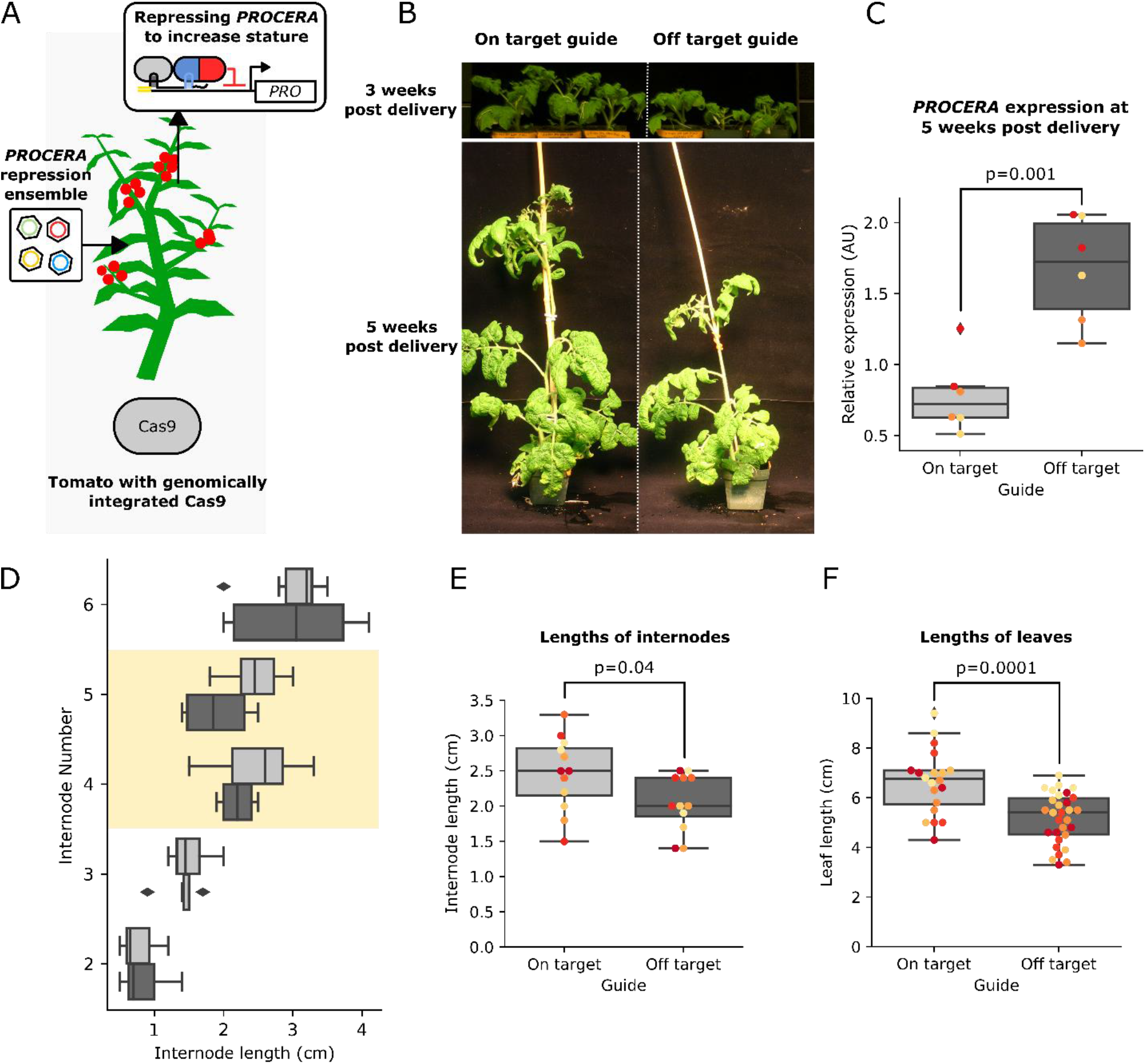
ViN 2.0 ensembles can repress *PROCERA* expression systemically and rapidly create increased stature and organ size. A) Schematic describing the ViN 2.0 repressor ensemble used to deliver a TPLN188 repressor and sgRNA scaffolds to target the *PROCERA* gene in a tomato line that stably expresses Cas9. The inset panels show the resultant reconstituted transcription factor at the *PROCERA* promoter. B) Representative images of plants treated with ensembles encoding off-target (left) and on-target (right) sgRNA scaffolds, 3 weeks (top) and 5 weeks (bottom) post vector delivery. C) Boxplot summarizing relative expression of *PROCERA*, normalized to the housekeeping gene *qCAC*, from systemic leaves of the plant lines described in panel A that were treated with ViN vectors encoding off-target (light grey) or on-target (dark grey) sgRNAs. Each dot of a different color represents data from independent biological replicates (n=3 per treatment). Reported p-values were calculated using a t-test. D) Boxplots summarizing the internode lengths of the plants described in panel A treated with ViN vectors encoding off-target (light grey) or on-target (dark grey) sgRNAs. Internode number 1 is at the bottom of the plant. The yellow region highlights the internodes in which we see a significant difference in size between plants treated with on-target and off-target sgRNAs. E) Boxplots summarizing the lengths of the fourth and fifth internodes, which are highlighted in yellow in D, of the plants described in panel A treated with ViN vectors encoding off-target (light grey) or on-target (dark grey) sgRNAs. Each dot of a different color represents data from an independent biological replicate (n=6 per treatment). Reported p-values were calculated using a t-test. F) Boxplots summarizing the lengths of leaves on the 8^th^ systemic branch of the plants described in panel A treated with ViN vectors encoding off-target (light grey) or on-target (dark grey) sgRNAs. Each dot of a different color represents data from an independent biological replicate (n=5 per treatment). Reported p-values were calculated using a t-test.

We delivered the ensemble targeting a repressor to *PROCERA* in parallel with another ensemble encoding off-target controls in Cas9-expressing tomato plants (var. M82) at the two true leaf stage and analyzed *PROCERA* expression in systemic leaves after five weeks. We observed a significant repression of *PROCERA* expression in plants treated with ensembles that encoded on-target sgRNAs compared to off-target controls (Figure 5B). We also observed the expected gradient of vector concentration decreasing from the point of delivery, demonstrating effective systemic movement of the vectors (Figure 5 Supplementary Figure 1).

The previously mentioned model of GA signaling^13^ and previous studies^29–33^ predict that repressing *PROCERA* expression would result in an increased stature and organ size. We phenotyped the internode length and observed a significant increase in the lengths of the fourth and fifth internodes (Figure 5C). We hypothesize that we do not see increased length in the first few internodes, as it takes time for the vector concentration and co-localization to build to the levels that can create physiologically relevant changes in the expression of *PROCERA*. We believe that the top internode was early in its development and not fully extended, which is why we did not see a significant difference there. This mirrors the pattern of change in leaf size we report in our experiments with ViN 1.0 in *N. benthamiana* in Figure 1. This pattern was further replicated in a second trial with independently grown plants, further confirming these results were not due to growth condistions or positional effects (Figure 5 Supplemental Figure 2B).

We also observed a significant increase in the length of systemic leaves in plants treated with ensembles encoding on-target sgRNAs compared to controls (Figure 5D). This is consistent with the repression of *DELLA* expression leading to increased tissue elongation^16^. These phenotypes were achieved within five weeks of vector delivery, compared to the months to years it would take to generate comparable phenotypes via transgenic approaches^30^. Taken together these results demonstrate how ViN 2.0 ensembles can be deployed in tomato to rapidly create transcriptional alterations for *in planta* hypothesis testing. They also show how these alterations can be guided by mechanistic models to rapidly engineer agronomically important traits in crops.

**Figure 5 Supplementary Figure 1.**
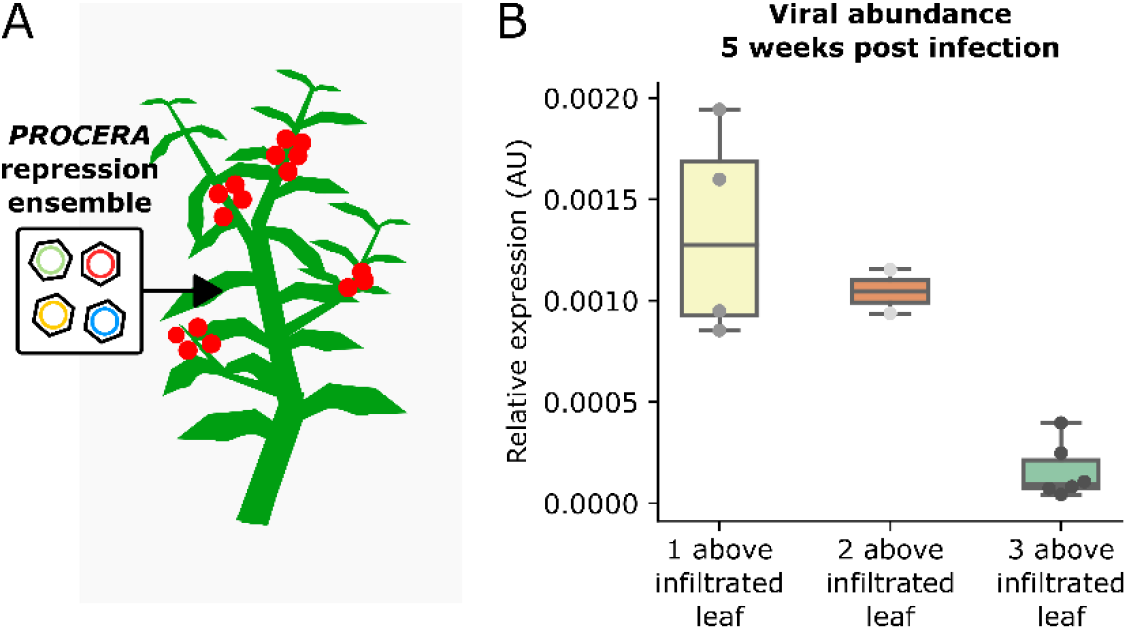
vector abundance decreases in progressively distal tissues. A) Schematic describing the ViN 2.0 repressor ensemble used to deliver a TPLN188 repressor and sgRNA scaffolds to target the *PROCERA* gene in a tomato line that stably expresses Cas9. B) Boxplot summarizing normalized levels of RNA of ViN vectors from systemic leaves of the plant lines described in panel A. Each dot represents data from independent biological replicates.

**Figure 5 Supplementary Figure 2.**
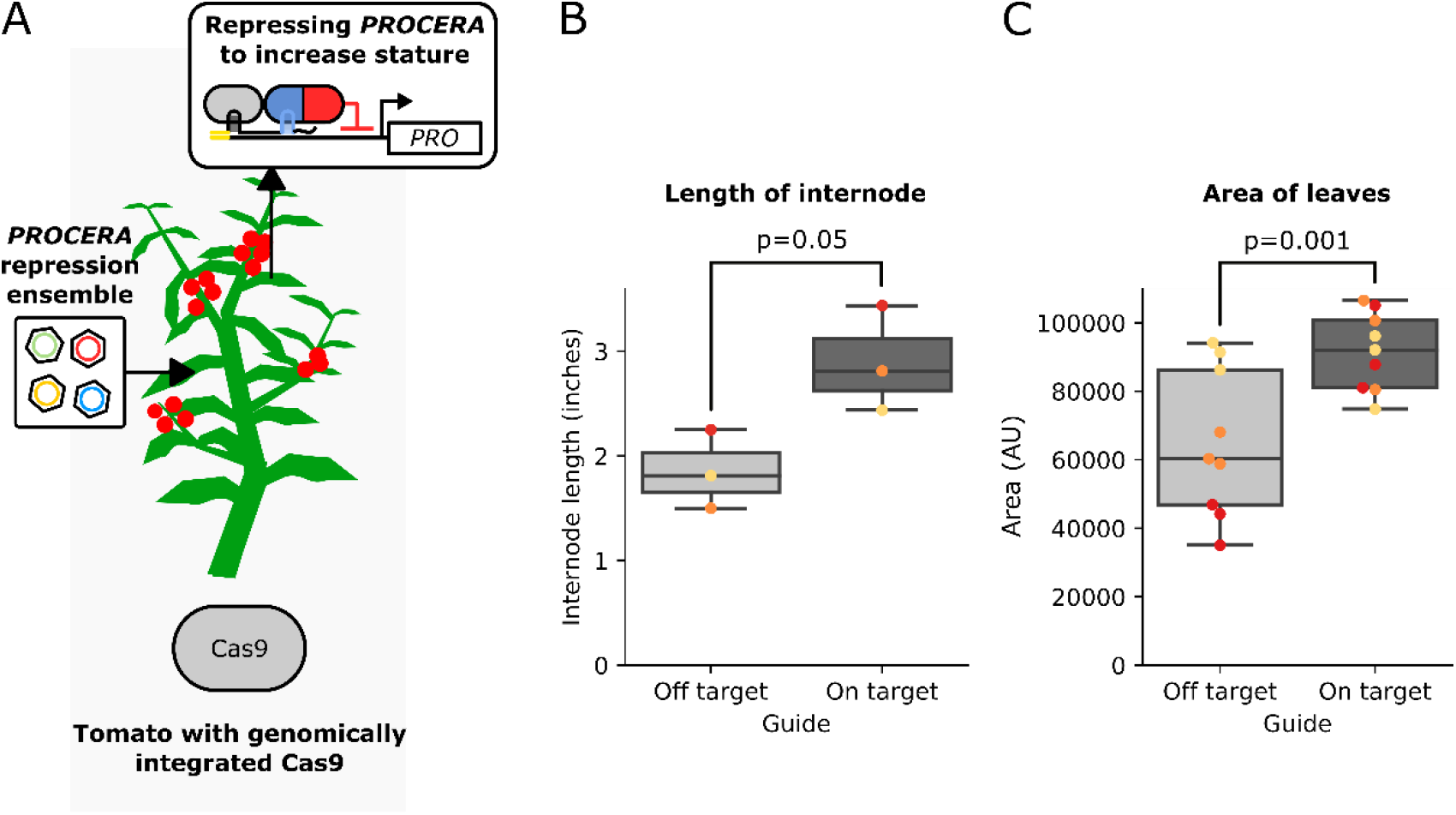
Phenotypic effects of ViN 2.0 ensembles repressing PROCERA can be replicated in a second set of independently grown plants. A) Schematic describing the ViN 2.0 repressor ensemble used to deliver a TPLN188 repressor and sgRNA scaffolds to target it to the *PROCERA* gene in a tomato line that stably expresses Cas9. B) Boxplots summarizing the lengths of the fourth internode of the plants described in panel A treated with ViN vectors encoding off-target (light grey) or on-target (dark grey) sgRNAs. Each dot of a different color represents data from an independent biological replicate (n=3 per treatment). Reported p-values were calculated using a t-test. C) Boxplots summarizing the area of leaves of the plants described in panel A treated with ViN vectors encoding off-target (light grey) or on-target (dark grey) sgRNAs. Each dot of the same color represents data from an independent biological replicate (n=3 per treatment). Reported p-values were calculated using a t-test.

## Discussion

In this work, we demonstrate how the ViN 1.0 platform can be used to rapidly reprogram both phytohormone-mediated and static transcriptional regulation in the model plants, *A. thaliana* and *N. benthamiana*. This expression alteration is observed across the plant and persists for several weeks. We show how ViN vectors can be deployed in HACR lines, guided by a model of GA signaling^13,16^, to alter the expression of key genes in this pathway and create predictable GA biosynthesis and dwarfing phenotypes. We also show how plant lines engineered to enable scaffold-based reconstitution of transcription factors allow simultaneous activation and repression of genes to tune multiple pathways and thereby stack traits, namely dwarfing and anthocyanin accumulation. These synthetic transcription factors have been shown capable of tuning gene expression^7,18^, making them powerful tools for plant engineering. Our results highlight how these tools might be used to reprogram the expression of multiple genes simultaneously to explore and engineer multigenic phenotypes. While we do observe some variability in the strength of regulation across biological replicates, the magnitude is comparable to variability observed in previously characterized transgenic lines stably expressing synthetic transcription factors^5^. The modest changes in gene expression we report, and the inability to create observable changes in expression for some genes, highlight a need for further development. In the future larger fold changes might be achieved by using different transcriptional effector domains or through recruitment of multiple effectors per sgRNA. Further expanding the existing toolbox of effectors encoded by ViN vectors to include more activators, repressors, and other enzymatic domains capable of epigenetic modifications are exciting avenues for future work that we are pursuing. The metabolic phenotypes we created serve as a proof-of-concept of how ViN vectors could be used to tune the expression of biosynthetic enzymes, like the *GA20ox* gene family, or transcription factors that regulate entire pathways, like *PAP1*, to rapidly engineer plant metabolism. VipariNama, when paired with high throughput phenotyping, will enable rapid screening of candidate genes at scale to elucidate the core components of poorly understood metabolic pathways.

We describe how VipariNama could be made even more flexible by using ViN 2.0 ensembles. According to our results, the incorporation of a movement enhancement motif is essential for the efficient functioning of this system. We show that this appears to be due to enhanced colocalization of the vectors, rather than enhanced concentration. In the future, single cell sequencing approaches could be used to further characterize the degree of co-localization of these vectors. There are several systemically mobile tRNA-like sequences that have been reported, some with tissue specific movement patterns^27^, and this represents a potential avenue to improve ViN vector performance or tissue specificity in the future.

While, ViN 2.0 does confer increased flexibility and obviates the need to build plant lines stably expressing transcription factor components, this approach does have some limitations. It can only be used to deploy relatively small transcriptional effectors due to the limited cargo capacity (<1kb) of these vectors. Additionally, while ViN 2.0 requires co-localization of at least two vectors in the same cell for effective regulation, ViN 1.0 does not. This might increase VIN 1.0’s efficacy, although we see no major evidence of this in the experiments reported in this work. A more thorough comparison using the same sgRNAs and effectors is needed to definitively determine their relative efficacy. Thanks to the broad host range of TRV^24^ and the fact that using ViN 2.0 only requires plants that stably express Cas9, these vectors can be rapidly and widely deployed, as Cas9 lines already exist for most major crops. This capability allowed us to deploy ViN 2.0 in a Cas9 expressing line of tomato and demonstrate systemic and persistent alteration of gene expression.

Previous studies of the phenotypic effects of perturbing the GA signaling pathway have focused on knockouts or overexpression of key genes^29–33^. These extreme cases, while mechanistically informative, do not generally represent reasonable agricultural interventions as they are often associated with pleiotropic effects. By utilizing synthetic transcription factors, we can make smaller changes and examine ranges of gene expression that might yield agriculturally beneficial phenotypes without the associated extreme pleiotropic effects. Our observation that *PAP1* activation, which was too subtle to be quantified experimentally, was able to reproducibly generate a measurable metabolic phenotype, highlights how small changes in gene expression can be phenotypically meaningful. Based on our results in *N. benthamiana*, tuning down the GA-induced expression of *GA20ox* genes is an avenue to reduce GA concentration in tissues and rapidly create dwarfing phenotypes. Our experiments in *A. thaliana* show another avenue to rapidly create dwarfing phenotypes is to statically repress the expression of the GA receptors, the *GID*s. Using ViN 2.0 we were able to extend this approach to tomato where we demonstrate that repressing the expression of the tomato *DELLA* protein, *PROCERA*, is an avenue to increase internode and leaf length. These results show, for the first time, that synthetic transcription factors can be applied to predictably increase or decrease plant size by modulating GA biosynthesis, GA perception, or GA-dependent regulation via tuning the expression of the *GA20ox*, *GID*, or *DELLA* genes respectively. The fact that we were able to use similar strategies across various plants points to this being a generic strategy to engineer plant size, an agriculturally relevant trait^4^. Being able to generically tune the GA signaling pathway has broad implications for improving the yield and geographical range of a wide range of mainstream crops like cotton and soybean, as well as orphan crops like teff and quinoa^4^.

The tomato phenotypes were generated within a few weeks of vector delivery, compared to the more than year it would take to generate similar phenotypes via transgenic interventions^30^. It was also achieved via an *Agrobacterium* infiltration, which requires significantly less time and technical skill than any transgenesis protocol. In the future we are excited to explore non-Agrobacterium based nucleic acid delivery methods, such as carbon nanotubes^34^, to expand these tools to species that cannot be easily infected by Agrobacterium. The capacity to avoid transgenesis also enables some common issues associated with the generation of stable lines to be side stepped. For example, the generational silencing of transgenes, the requirement of strong and constitutive promoters, and the necessity to screen lines to account for variation associated with the genomic context of transgene insertion. However, ViN vectors do introduce some new sources of variation associated with viral movement, tissue tropism, and potential viral pleiotropic effects, which will have to be investigated further in future studies. Additionally, by obviating the cost and skill associated with transgenesis, these tools make plant synthetic biology more accessible to the wider plant biology community.

However, while ViN vectors are powerful new tools, there are many areas for further improvement. An important area is engineering an effective and genetically stable containment and clearing system, in case they somehow escape the contained lab conditions we use them in. One potential strategy we are exploring is incorporating chemically cleavable motifs, called aptazymes^35^, into the viral genome in conserved regions via synonymous mutations. Another is removal of structural viral proteins critical for vector transmission^24^. However, truly safe vectors will likely require multiple overlapping containment mechanisms that are independent of each other in the same vector to minimize the chances of escape. Engineering ViN vectors to be asymptomatic in all conditions is another area that requires work. While we did not see strong symptoms associated with viral infection in our experimental conditions, low temperatures (20-22°C) can result viral symptoms in *N. benthamiana*. We are developing high throughput strategies to rapidly engineer these and other aspects of viral behavior to make them more robust to environmental changes.

The results presented here highlight how VipariNama accelerates synthetic transcription factor-mediated hypothesis testing by obviating transgenesis. This is the first step to building predictive models that could be used for precision forward engineering. VipariNama will also allow plant engineers to rapidly iterate through different targets and effectors to identify optimal synthetic transcription factor mediated interventions for crop improvement. We hope these tools will empower the plant science community to deliver solutions to the pressing issues facing global agriculture.

## Materials and methods

### Transgenic line generation

The transgenic GA-HACR expressing *N. benthamiana* lines used in this paper were generated by transforming the previously published GA-HACR^5^ plasmid into wildtype *N. benthamiana*. This construct includes expression cassettes for the GA-HACR itself, which is a dCas9 protein fused to the *A. thaliana* DELLA protein *RGA* and a truncation of the *A. thaliana* repressor TOPLESS, as well as for a venus-luciferase fusion reporter and a sgRNA that targets the GA-HACR to this reporter. The transgenesis protocol previously published by Sparkes *et al*^36^ was used to generate stable transgenic lines and a highly expressing T2 line was used to perform all the experiments in this paper.

The *A. thaliana* transgenic lines described in this paper were generated by integrating either the HACKER locus 3 or HACKER locus 20 construct into the genome via floral dip^37^. Both HACKER loci are tDNAs that contain expression cassettes for the constitutive expression of a nuclease active Cas9 codon optimized for expression in *A. thaliana*. They also both contain an expression cassette for the expression of a COM RNA binding protein fused to a VP64 activator domain via a flexible linker. The HACKER locus 3 contains an additional expression cassette that constitutively expresses a PCP RNA binding domain fused to a truncation of the *A. thaliana* repressor *TOPLESS*. This polypeptide is fused, via a P2A peptide, to an MCP RNA binding domain which is in turn fused to an auxin degron and a truncation of *TOPLESS*. This allows these two RNA binding protein effector fusions to be post-translationally separated, theoretically enabling simultaneous repression, as well as well as hormone regulated repression. Finally, the HACKER 3 locus also encodes a Venus reporter, whereas the HACKER 20 locus encodes both a *Renilla* luciferase and a firefly luciferase reporter. These constructs were built using a two-step Goldengate assembly^38^, with some parts from the previously published genome engineering kit^39^.

The Cas9 expressing lines of both *N. benthamiana* and *S. lycopersicum* (M82) used in this work were generated previously^40^. A list of all the plasmids described is available in Table 1 and the associated genbank sequence files are available in the supplementary information.

**Table 1 -.** List of plasmids.

| Plasmid Number | Plasmid description |
| --- | --- |
| P38 | HACKER locus 3 |
| p145 | PHD6 (p2301Y-tOCS-pUBQ1:NLS-Venus-LucPlus-tUBQ1-pU6:pUBQ1_gRNA_Target1-tU6-pUBQ10:dCas9-RGA1-TPLRD2-tNos) |
| P178 | HACKER locus 20 |
| P254 | TRV2-NLS-PCP-SRDX-STOP |
| P262 | TRV2-NLS-PCP-SRDX-STOP-Ft_peptide |
| P263 | TRV2-NLS-COM-VP64-STOP-Ft_peptide |
| P266 | TRV2-NbDFR_Target2_truncated-gRNAhandle-COM-Ft_peptide |
| P267 | TRV2-NbDFR_Target3_truncated-gRNAhandle-COM-Ft_peptide |
| P269 | TRV2-NbPDS3.1_truncated_target_2_gRNA-PP7 |
| P270 | TRV2-NbPDS3.1_truncated_target_2_gRNA-PP7-Ft_peptide |
| P272 | TRV2-NbPDS3.2_truncated_target_1_gRNA-PP7-Ft_peptide |
| P295 | TRV2-NLS-PCP-TPL_N188-STOP-Ft_peptide |
| P309 | TRV2-NbPDS3.1_truncated_target_1_gRNA-PP7 |
| P310 | TRV2-NbPDS3.1_truncated_target_1_gRNA-PP7-Ft_peptide |
| P312 | TRV2-NbPDS3.2_truncated_target_2_gRNA-PP7-Ft_peptide |
| P383 | TRV2_pAtGID1a_target_3_truncated_gRNA-PP7-Ft_peptide |
| P384 | TRV2_pAtGID1b_target_3_truncated_gRNA-PP7-Ft_peptide |
| P385 | TRV2_pAtGID1c_target_3_truncated_gRNA-PP7-Ft_peptide |
| P386 | TRV2-AtPAP1_truncated_target_1_gRNA-COM-Ft_peptide |
| P387 | TRV2-AtPAP1_truncated_target_2_gRNA-COM-Ft_peptide |
| P388 | TRV2-AtPAP1_truncated_target_3_gRNA-COM-Ft_peptide |
| P389 | TRV2-pNbGA20ox1-1_truncated_target_1_gRNA-PP7-Ft_peptide |
| P390 | TRV2-pNbGA20ox1-1_truncated_target_2_gRNA-PP7-Ft_peptide |
| P391 | TRV2-pNbGA20ox1-2_truncated_target_1_gRNA-PP7-Ft_peptide |
| P392 | TRV2-pNbGA20ox1-2_truncated_target_2_gRNA-PP7-Ft_peptide |
| P393 | TRV2-pNbGA20ox1-3_truncated_target_1_gRNA-PP7-Ft_peptide |
| P394 | TRV2-pNbGA20ox1-3_truncated_target_2_gRNA-PP7-Ft_peptide |
| P395 | TRV2-pNbGA20ox1d-1_truncated_target_1_gRNA-PP7-Ft_peptide |
| P396 | TRV2-pNbGA20ox1d-1_truncated_target_2_gRNA-PP7-Ft_peptide |
| P397 | TRV2-pNbGA20ox1d-2_truncated_target_1_gRNA-PP7-Ft_peptide |
| P398 | TRV2-pNbGA20ox1d-2_truncated_target_2_gRNA-PP7-Ft_peptide |
| P472 | TRV2_SIProcera_Target_3_truncated_gRNA-2xPP7-FT_peptide |
| P473 | TRV2_SIProcera_Target_4_truncated_gRNA-2xPP7-FT_peptide |
| P474 | TRV2_SIProcera_Target_5_truncated_gRNA-2xPP7-FT_peptide |

### ViN vector construction

The ViN vectors described in this work were all modified versions of the TRV2 genome^41^. These add an additional sub-genomic promoter and the RNA we which to deliver downstream of the coat protein coding sequence. The ViN vectors that encode a truncated guide RNA scaffold consist of a guide RNA with a 14 base pair long target sequence, the handle sequence, and the motifs which interact specifically with either the PCP or COM RNA binding proteins, depending on the experiment. All the sequences used for on-target and off-target guide RNA scaffolds are listed in Table 2.

**Table 2 –.**
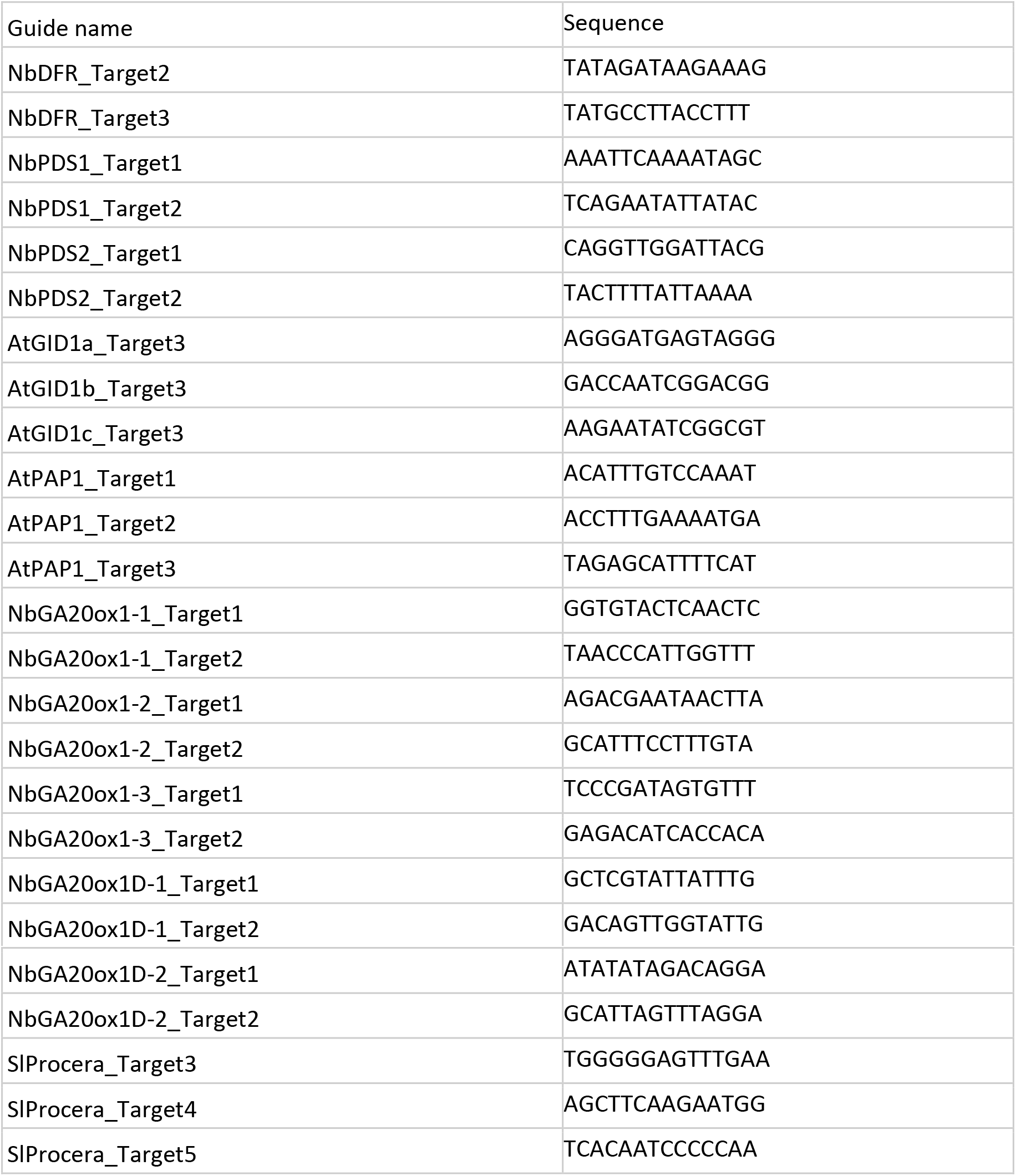
List of guide sequences.

The ViN vectors used in ViN 2.0 ensembles to encode effectors contain coding sequences for either the SRDX repressor domain or the 188 N-terminal residues of *TOPLESS* fused to a PCP RNA binding domain, or a VP64 activator domain fused to the COM RNA binding domain. All the vectors, unless specified, encode the tRNA-like sequence in first 102 base pairs of the *A. thaliana* FLOWERING LOCUS T transcript 3’ of the cargo as a movement enhancement sequence. All these vectors were built using a one-step Goldengate assembly.

### Agrobacterium-based vector delivery

In all experiments ViN vectors were delivered via *Agrobacterium* infiltration of leaves of young plants^36^. Individual *Agrobacterium* (GV3101) containing the tDNAs encoding the ViN vectors or the TRV1 genome were cultured overnight in LB media with Kanamycin (50 mg/ml) and Gentamycin (50 mg/ml) selection on shaker at 220rmp and 28°C. The next day (18-20 hours later), once the cultures were at confluent growth, they were spun down at 2500xg for 10 minutes and washed twice with infiltration media (10 mM MgCl2, 10 mM 2-(N-morpholino) ethanesufonic acid (MES) at pH 5.6). For infiltrations into *N. benthamiana* these cultures were then resuspended in infiltration media at an OD600 of 0.6 and allowed to rest room temperature for 2-3 hours prior to infiltration. For infiltrations into *A. thaliana* these cultures were then resuspended in infiltration media with Acetosyringone (200uM final concentration) at an OD600 of 1 and allowed to rest room temperature for 2-3 hours prior to infiltration. For infiltrations into *S. lycopersicum* these cultures were then resuspended in infiltration media with Acetosyringone (200uM final concentration) at an OD600 of 2 and allowed to rest room temperature for 2-3 hours prior to infiltration. For all assays performed a 1:1 ratio of by volume of *Agrobacterium* containing TRV1 to TRV2 was used. For ViN 1.0 the final infiltrated mixtures of *Agrobacterium* contained equal volumes of strains that contained the sgRNA encoding TRV2s. For ViN 2.0 the final infiltrated mixtures of *Agrobacterium* contained equal volumes of strains that contained the sgRNA encoding TRV2s as well as one volume of the strain that contained an RNA binding protein-effector fusion encoding TRV2 per volume of the sgRNA encoding TRV2s.

### GA-HACR-based luciferase assay

For the luciferase assays that were used as a proxy for GA concentration with the GA-HACR *N. benthamiana* lines treated with ViN vectors, systemic leaves were assayed at the time of phenotyping, 9 weeks post vector delivery. Luciferin (100uM final concentration in water) was infiltrated into the leaves being assayed and they were removed from the plants and imaged after five minutes. The leaves were imaged using a CCD camera-based luciferase imaging eight-minute exposures of plant tissue were taken in using a UVP BioImaging Systems EpiChemi3 Darkroom with a 10-minute exposure. ImageJ^42^ was used to quantify the luminescence in each infiltrated zone with three technical replicates per measurement.

### Expression analysis via qRT-PCR

For all expression analysis in this work, tissue was collected from systemic tissues on plants treated with ViN vectors and RNA was extracted using the TRIZOL reagent (Invitrogen). After a DNAse treatment with the TURBO-DNAse kit (Invitrogen) the concentration of RNA of the targeted genes, the ViN vectors and housekeeping genes was quantified using qRT-PCR performed with the one-step SuperScript RT-PCR kit (Invitrogen) on a BioRAD thermocycler. Reactions were scaled down to 12.5ul with 50ng of RNA per reaction to conserve reagents. Between one and two technical replicates were performed on all samples. The qPCR primers used are listed in Table 3.

**Table 3 –.**
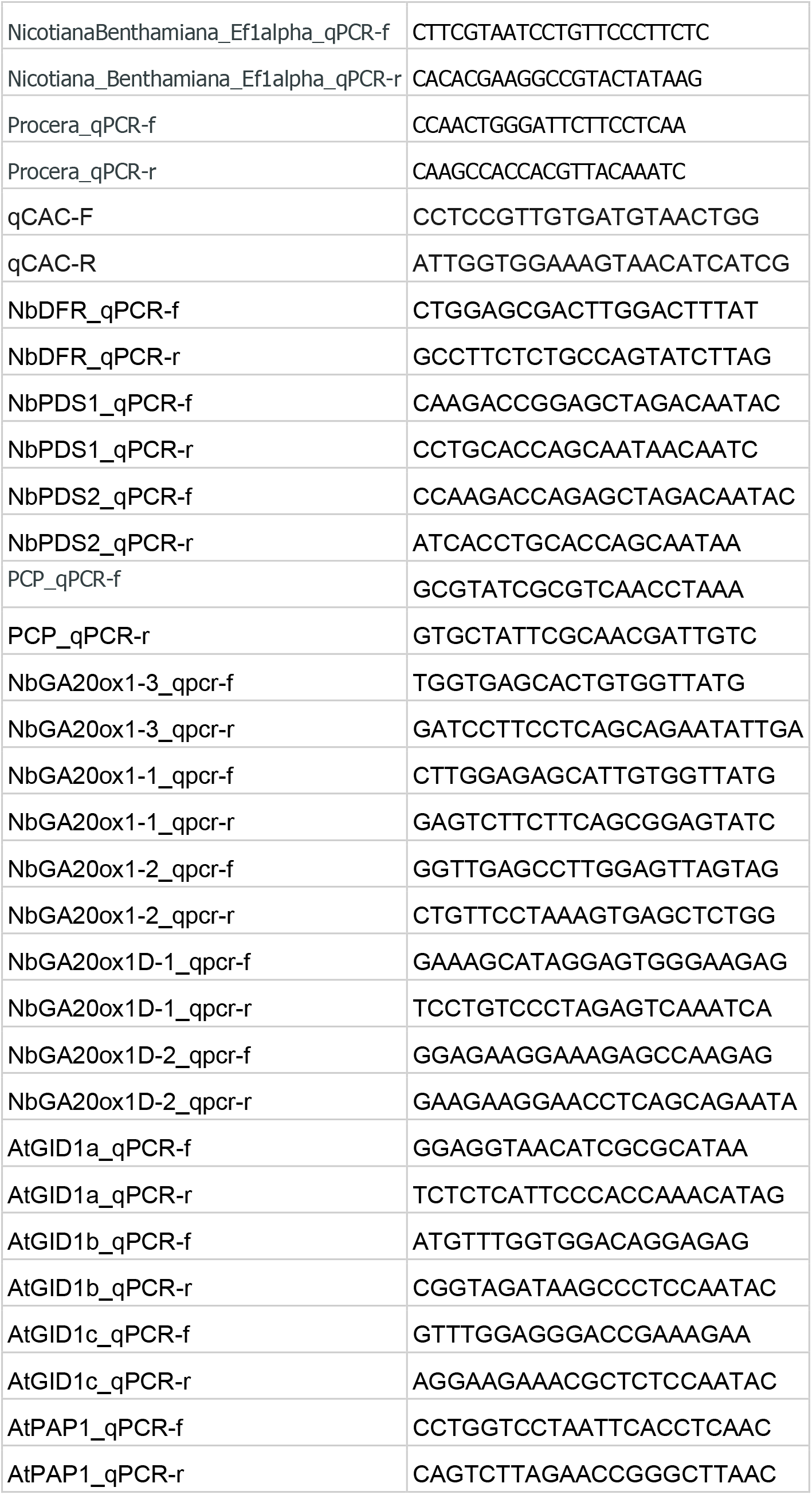
List of qPCR primers.

### Phenotyping *N. benthamiana* leaf size

The GA-HACR expressing *N. benthamiana* plants were treated with ViN vectors and grown in a Conviron chamber set to a 18 hour day (26°C) and 6 hour night (24°C) light cycle. The temperature conditions were observed to be critical to prevent viral necrosis, if the temperature was dropped too low. The plants were placed in a well-mixed pattern across treatments in the chamber to prevent location specific effects confounding the results. These plants were phenotyped nine weeks post vector delivery. The leaves on the main stem were removed and sequentially arrayed on a uniformly lit stage with a fixed camera distance and photographed. The leaf size was then calculated from these images using ImageJ^42^. The raw images used to make measurements are included in the supplementary data.

### Phenotyping Arabidopsis rosette leaf size and color

The *A. thaliana* plant lines were treated with ViN vectors and grown in a 18 hour day and 6 hour night light cycle at 22°C. These plants were phenotyped three weeks post vector delivery. The inflorescences were removed, and the plants were mounted on a uniformly lit stage and imaged with a Canon DSLR mounted on a fixed tripod to maintain consistent focal distance. To estimate the anthocyanin concentration in the rosette leaves the images were then analyzed in imageJ and the mean Blue and green signals from three different rosette leaves per plant were recorded. These were then used to calculate the Blue to green ratio, which has been established as a proxy that correlates with anthocyanin accumulation^23^. To phenotype the size of the rosette leaves, the rosettes were dissected and placed sequentially on a scanner and imaged. These scans were then analyzed in ImageJ and the size of the individual leaves was recorded. The raw images that were analyzed are available in the supplementary data.

### Phenotyping tomato leaf and internode length

The Cas9 expressing tomato (M82) lines were treated with ViN vectors and grown in a Conviron chamber set to a 18 hour day (26°C) and 6 hour night (24°C) light cycle. The plants were placed in a well-mixed pattern across treatments in the chamber to prevent location specific effects confounding the results. These plants were phenotyped fifty days post viral delivery. The length of the internodes and the leaves were physically measured using a measuring tape. Images of the plants were also taken for one set of plants over the course of their growth, as presented in Figure 5.

### Data analysis and plotting

All the data analysis was performed in python and the associated jupyter notebooks are available in the supplementary data. All p values reported were calculated using the t-test function in the scipy package. All the data presented was plotted using the seaborn package^43^ in python.

---

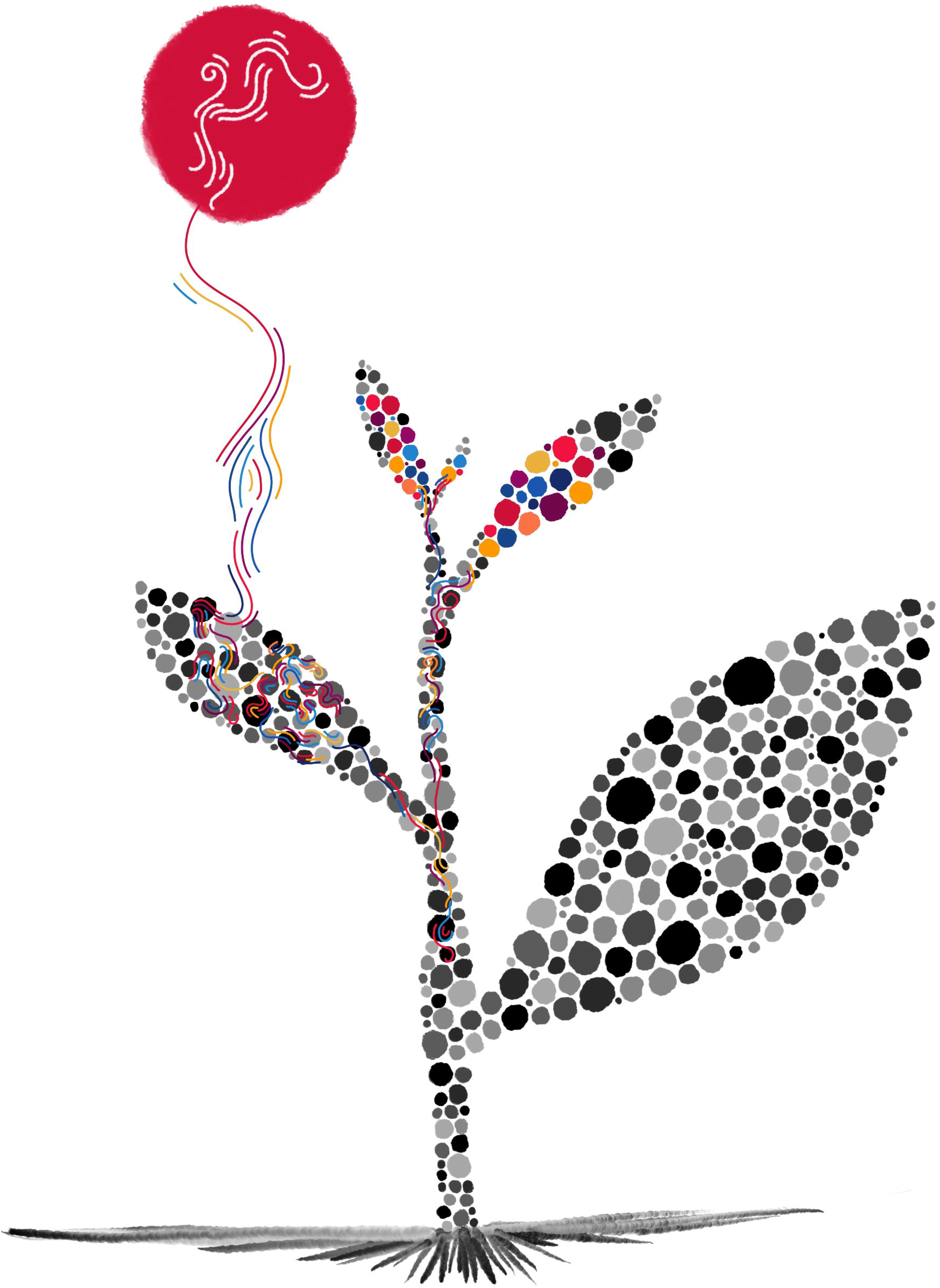

